# Gene regulatory network of human GM-CSF secreting T helper cells

**DOI:** 10.1101/555433

**Authors:** Szabolcs Éliás, Angelika Schmidt, David Gomez-Cabrero, Jesper Tegnér

## Abstract

GM-CSF produced by autoreactive CD4 positive T helper cells is involved in the pathogenesis of autoimmune diseases, such as Multiple Sclerosis. However, the molecular regulators that establish and maintain the features of GM-CSF positive CD4 T cells are unknown. In order to identify these regulators, we isolated human GM-CSF producing CD4 T cells from human peripheral blood by using a cytokine capture assay. We compared these cells to the corresponding GM-CSF negative fraction, and furthermore, we studied naïve CD4 T cells, memory CD4 T cells and bulk CD4 T cells from the same individuals as additional control cell populations. As a result, we provide a rich resource of integrated chromatin accessibility (ATAC-seq) and transcriptome (RNA-seq) data from these primary human CD4 T cell subsets, and we show that the identified signatures are associated with human autoimmune disease, especially Multiple Sclerosis. By combining information about mRNA expression, DNA accessibility and predicted transcription factor binding, we reconstructed directed gene regulatory networks connecting transcription factors to their targets, which comprise putative key regulators of human GM-CSF positive CD4 T cells as well as memory CD4 T cells. Our results suggest potential therapeutic targets to be investigated in the future in human autoimmune disease.

## INTRODUCTION

CD4 positive T helper cells (Th) are crucial players in the immune system which exert their effects mainly by producing cytokines. CD4 T cell subsets are usually classified based on expression of “lineage-defining” transcription factors (TFs) as well as the signature cytokines they secrete [1]. However, the distinction is not clear-cut, since different signature cytokines can be expressed simultaneously and plasticity between subsets occurs [2,3]. In view of the “classical” distinction of CD4 T cell subsets, particularly Th1 and Th17 subsets are involved in the establishment of autoimmune diseases such as Multiple Sclerosis (MS) and the corresponding rodent model Experimental Autoimmune Encephalomyelitis (EAE), which are thought to be driven by the T cell-released cytokines Interleukin-17 (IL-17), Interferon-γ (IFN-γ), IL-22 and granulocyte-macrophage colony-stimulating factor (GM-CSF). Of these, GM-CSF was determined as the key cytokine in EAE pathogenesis, since only knocking out GM-CSF (but neither IFN-γ, IL-17A nor IL-17F) could completely protect the animals from induced EAE [4–6]. Furthermore, it has been demonstrated that specifically the GM-CSF produced by autoreactive T cells was necessary for EAE induction, while T cell-produced IFN-γ and IL-17 were dispensable [7–9]. In line with these results, GM-CSF expression by Th cells was required for neuroinflammation in EAE, and even in the presence of IFN-γ and IL-17A producing Th cells, pathogenicity was lost upon loss of GM-CSF [10]. Importantly, in humans the fraction of GM-CSF positive (and IFN-γ positive) cells within CD4 T cells was elevated in MS patients’ cerebrospinal fluid compared to controls, while IL-17A positive cell fractions were not strikingly different in these reports [11,12]. The fraction of GM-CSF positive and IFN-γ positive cells was also increased in peripheral blood of MS patients in one report [13], but not in another [11]. Similarly, enhanced fractions of GM-CSF producing CD4 T cells have been observed in synovial fluid of patients with juvenile arthritis along with the well-known enhanced GM-CSF levels in synovial fluid [14,15]. Interestingly, an expanded Th subset producing GM-CSF was found in blood and CNS of MS patients and could be diminished by disease-modifying therapy [16], suggesting GM-CSF producing Th cells to be an attractive therapeutic target. Of note, targeting GM-CSF in MS or arthritis is subject to several ongoing clinical studies, highlighting the importance of this cytokine in these diseases [15]. Based on this cumulative evidence of the significance of GM-CSF producing CD4 T cells in human autoimmune disease, understanding the factors driving and defining GM-CSF positive T cells would be of utmost importance for targeting them therapeutically.

Beyond their established pathogenic role in autoimmune diseases, GM-CSF producing Th cells have also been implicated in other, inflammatory diseases. In sepsis, enhanced fractions of GM-CSF producing T cells were associated with a poor outcome [17]. Notably, GM-CSF producing Th cells have also been implicated in SARS-CoV-2 infection, especially in patients with severe course of coronavirus disease 2019 (COVID-19) [18–20]. Due to the pleiotropic roles of GM-CSF in immune disease and lung inflammation, GM-CSF-targeting therapeutic approaches are currently explored in clinical trials to treat COVID-19 [21]. Murine CD4 T cell populations expressing IL-17A and GM-CSF have been observed and termed “pathogenic Th17” cells because they have the potential to induce EAE [22–24]. However, only one of these studies showed co-expression of both cytokines on the single cell level [24]. Although a T cell can express GM-CSF simultaneously with other cytokines such as IFN-γ, a “GM-CSF only” producing murine T cell subset was also proposed and associated with enhanced encephalitogenic activity over IL-17 and IFN-γ producing T cells in EAE [9]. The existence of a corresponding separate “GM-CSF-only” human T cell subset has also been proposed [25], because a substantial subset of human GM-CSF positive CD4 T cells produces GM-CSF in the absence of any other classical Th lineage-defining cytokines, transcription factors or surface markers [11,26]. Furthermore, GM-CSF producing CD4 T cells are induced by different sets of cytokines compared to other Th cell subsets [25,27]. In fact, GM-CSF and IL-17A expression by human CD4 T cells has been found to be mutually exclusive on single cell level [11] or at least less frequent than co-expression of IFN-γ and GM-CSF [26,27].

Despite the importance of GM-CSF producing T cells, there is no specific marker to distinguish such cells from others to date. Although combinations of presence and absence of non-exclusive surface markers has been useful to delineate “GM-CSF-only” cells [11], the fraction of GM-CSF producing cells that also produces other cytokines such as IFN-γ is excluded by this approach. Furthermore, the driving molecules for GM-CSF production remain unclear. Together, these observations suggest that the characterization of human GM-CSF positive CD4 T cells isolated based on their functional profile (GM-CSF production) rather than by distinction of the “classical” Th1 and Th17-like phenotypic markers may enable the identification of factors regulating GM-CSF production in CD4 T cells. Hence, in order to understand the regulators and molecular patterns defining human GM-CSF positive CD4 T cells, we here isolated those cells actively secreting GM-CSF from human peripheral blood *ex vivo* by cytokine “capture” assay, starting with bulk CD4 T cells. We then studied their transcriptome by RNA-sequencing (RNA-seq). Studying a single data type like mRNA expression separately may not be sufficient for identification of all regulatory factors, since TFs themselves are often regulated by post-transcriptional modifications, intracellular translocation or co-binding with other TFs, rather than by changes in their gene expression. Thus, we assessed in parallel the DNA accessibility of the same samples in order to gain a global picture of putative TF binding patterns and enabling integration with the expression of regulated target genes on RNA level. Due to the limited number of primary, *ex vivo* isolated GM-CSF positive human cells, we employed a recently described highly sensitive method, assay for transposase-accessible chromatin using sequencing (ATAC-seq) [28] to study DNA accessibility from 50,000 cells per sample. As a control, we used the respective GM-CSF depleted (“GM-CSF negative”) fraction derived from the capture assay procedure. Since GM-CSF positive cells may differ from GM-CSF negative CD4 T cells simply by containing largely reduced fractions of naïve cells, we furthermore undertook RNA-seq and ATAC-seq profiling of several control cell populations that is, naïve CD4 T cells and memory CD4 T cells. As an additional control, we studied bulk CD4 T cells without capture assay procedure.

This study hence reveals molecular patterns specific for GM-CSF positive CD4 T cells or shared with memory, naïve or bulk CD4 cells. To our knowledge, this is the first study of global molecular signatures of GM-CSF positive CD4 T cells derived *ex vivo* without restimulation. Besides serving as a control we furthermore provide a novel resource of ATAC-seq and RNA-seq data of human primary naïve, memory and bulk CD4 T cells from several human healthy donors. A large body of knowledge exists on molecular signatures and regulation of human naïve and memory T cells [29]. This encompasses large consortium efforts to map human memory and naïve CD4 T cell subsets’ transcriptomes and epigenomes including chromatin accessibility [30], albeit these authors did not use the ATAC-seq method. Recently, few reports using ATAC-seq for T cells have been published and the impactful results support the power of the methodology. Many of these studies focus on murine CD8 T cell differentiation and exhaustion [31–35], and several recent studies also comprise ATAC-seq on CD4 T cells [10,28,36–40]. However, none of these works studied all the types of CD4 T cell subsets we analyzed here.

Through interpreted, integrative analysis of mRNA expression and DNA accessibility data from primary human CD4 T cell subsets we provide novel gene regulatory networks underlying GM-CSF production as well as the memory phenotype in human CD4 T cells, and we propose novel key TFs regulating these cells. The enrichment of the identified genes for human immune system diseases and specifically MS for GM-CSF positive cells underlines the clinical relevance of our data, which may be exploited in a multitude of basic and applied immunology studies in the future.

## MATERIALS AND METHODS

### Ethics Statement

Peripheral blood mononuclear cells were freshly isolated from anonymized healthy donor buffy coats purchased from the Karolinska University Hospital (Karolinska Universitetssjukhuset, Huddinge), Sweden. Research was performed according to the national Swedish ethical regulations (ethical review act, SFS number 2003:460).

### Experimental methods

#### PBMC and T cell isolation

Human peripheral blood mononuclear cells (PBMCs) were isolated using Ficoll-Paque gradient centrifugation from buffy coats according to standard procedures. In brief, buffy coats were diluted in PBS, layered on Ficoll-Paque (GE healthcare) and centrifuged at 1200xg for 20 min without break. Subsequently, the PBMC ring was collected. PBMCs were washed with PBS (450xg, 10 min) and monocytes were depleted by plastic adherence in RPMI 1640 medium containing GlutaMAX (LifeTechnologies, Thermo Fisher Scientific) and 10% (v/v) heat inactivated fetal bovine serum (FBS; Gibco Performance Plus certified; Thermo Fisher Scientific) for 60-80 minutes. Platelets were removed by centrifugation (200xg, 5-10 min, 20°C; 4 times). Subsequently, human naïve, memory, and total CD4 T cells were isolated negatively (“untouched”) in parallel by magnetic activated cell sorting (MACS) from each donor. The following MACS kits were used according to the instructions from the manufacturer (Miltenyi Biotec): human naïve CD4^+^ T cell isolation kit II, human memory CD4^+^ T cell isolation kit II, and human CD4^+^ T cell isolation kit II. The purity of naïve, memory, and total CD4 T cells was controlled by flow cytometry (see below). Cells were counted with the Countess Automated Cell Counter (Invitrogen) and viability (determined by Trypan blue stain) was 96.5±1.5% (mean±SD). T cells were cultured at 37°C and 5% CO_2_ in serum-free X-Vivo 15 medium (Lonza) supplemented with GlutaMAX (Gibco), unless otherwise stated. T cells were rested overnight before sample preparation for RNA-seq, ATAC-seq, and GM-CSF secretion assay (see below). 6 healthy male anonymized donors (age 35.7±8.5 years, mean±SD; range 22-44 years) were used for molecular profiling.

#### GM-CSF secretion assay (“capture assay”)

Total CD4 T cells were isolated and rested as above, before “capturing” GM-CSF producing cells using the GM-CSF Secretion Assay - Cell Enrichment and Detection Kit (PE), human (Miltenyi Biotec) according to the manufacturer’s instructions with the following modifications and details. 65-150×10^6^ (98.3×10^6^±33.7×10^6^, mean±SD) unstimulated purified CD4 T cells were centrifuged (300xg, 10 min), X-Vivo 15 medium was removed completely, and cells were washed with 15 ml MACS buffer (0.5 % human serum albumin (HSA) and 2 mM EDTA in PBS, 4°C). Cells were resuspended in ice-cold RPMI 1640 medium including GlutaMAX (LifeTechnologies, Thermo Fisher Scientific) and 10% (v/v) FBS (Gibco Performance Plus certified, heat inactivated; Thermo Fisher Scientific). GM-CSF catch reagent was added, mixed, and incubated for 5 minutes on ice. The GM-CSF secretion period was performed by adding pre-warmed (37°C) RPMI 1640 medium (containing GlutaMAX and 10% FBS) to a cell density of 1×10^6^ cells/ml, under continuous rotation (10 rpm orbital mixing) of the cells for 45 minutes at 37°C/5% CO_2_. Labeling cells with GM-CSF Detection Antibody (Biotin) and Anti-Biotin-PE, and magnetic labeling with Anti-PE MicroBeads UltraPure, were performed as per instructions and cells were washed in 50 ml MACS buffer. Cells were resuspended in 3 ml MACS buffer, cell suspensions were filtered with 30 μm Filcon strainer (BD Biosciences) and magnetic separation was performed on LS columns (Miltenyi Biotec) according to standard protocols. For Donor A, the GM-CSF+ eluate was passed over a second MS column following the LS column procedure, but since a second column did not increase purity but led to loss of cells (data not shown), for all other donors the GM-CSF+ eluate from the first LS column was used for subsequent analyses. The GM-CSF- fraction from the flow-through was passed over a second column (except for Donor A) to increase purity of the negative fraction. Yield of GM-CSF+ cells was 1.2±0.6% of CD4 T cells (mean±SD) and purity was controlled by flow cytometry (see below).

#### Flow cytometry

Purity of MACS-isolated naïve CD4 T cells, memory CD4 T cells and total CD4 T cells was verified by surface staining using the following antibodies (all against the human proteins): anti-CD4-PerCP (clone SK3, BD Biosciences), CD45RA-FITC (clone T6D11, Miltenyi Biotec), CD45RO-PE (clone UCHL1, BD Biosciences), CD19-APC (Clone HIB19, BD Biosciences), CD8-eFlour450 (clone OKT8, eBiosciences). Staining was performed in the dark with antibody dilutions in FACS buffer (PBS with 0.5% HSA) for 15 min at 20°C. Single stained PBMC or T cell samples were used as compensation controls. Cells were washed once with PBS, resuspended in FACS buffer and acquired on a CyAn ADP 9-Color Analyzer (Beckman Coulter). Compensation was performed with the CyAn software (Summit) tool. Purity of GM-CSF+ and GM-CSF- samples after GM-CSF secretion assay from bulk CD4 T cells was performed by measuring the fraction of PE-labeled (GM-CSF positive) cells in a CyAn ADP 9-Color Analyzer (Beckman Coulter). Cell purities and gating strategy are shown in Figure 1 and Supplementary Figure 1. Flow cytometry data analysis and visualization was performed using the FlowJo software v7.6.5 (Tree Star), and exported percentage values were plotted in GraphPad Prism v7.02 (GraphPad Software, Inc.).

**Figure 1.**
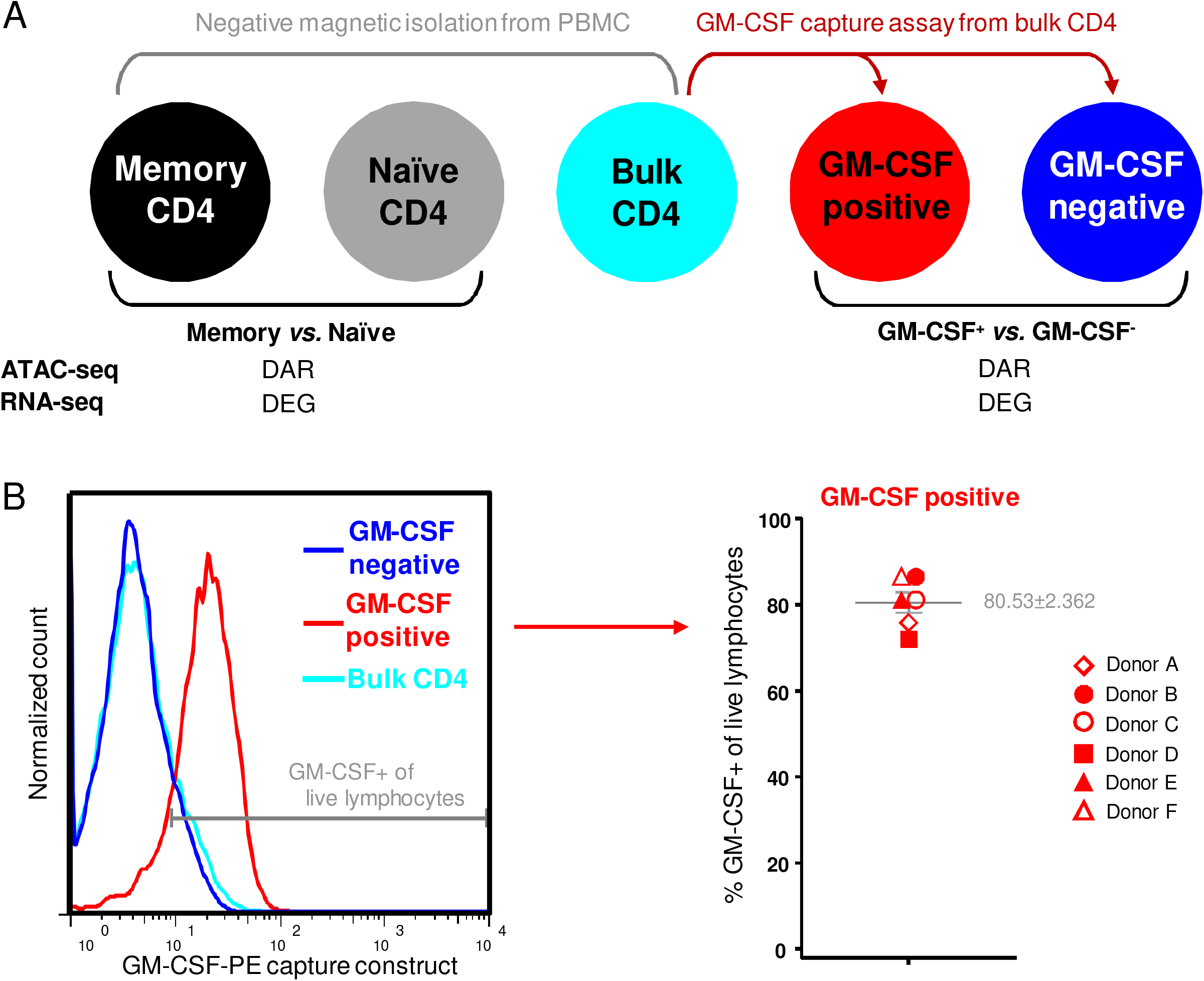
Experimental setup and quality control for human CD4 T cell transcriptome and chromatin accessibility analysis. (**A**) Human PBMCs for each donor were split in three fractions, and memory CD4 T cells, naïve CD4 T cells and bulk (total) CD4 T cells were isolated in parallel by negative (“untouched”) magnetic isolation (cell purities, see Supplementary Figure S1). From bulk CD4 T cells, GM-CSF-secreting cells (GM-CSF positive) were captured, and the negative fraction was used as corresponding GM-CSF negative population. The five indicated cell populations were used for molecular profiling by ATAC-seq and RNA-seq. Differentially accessible DNA regions (DAR) and differentially expressed genes (DEG) were determined for the comparison of memory *versus* naïve CD4 cells and for GM-CSF positive *versus* GM-CSF negative CD4 cells respectively. (**B**) High purity of isolated (captured) GM-CSF+ CD4 T cells was confirmed by flow cytometry. The histograms show the signal of the GM-CSF capture construct for the indicated cell populations, pre-gated on live singlet lymphocytes based on forward scatter, side scatter and pulse width. Here, bulk CD4 cells represent an aliquot taken after labelling with the GM-CSF-PE capture construct, but without further capture assay procedure. The left panel shows a representative donor and the right panel shows summarized data for n=6 donors from 4 independent experiments (mean and SEM are indicated in grey). Each donor is represented by a symbol; same symbol shape (but filled or unfilled) indicate donors processed within the same experiment.

#### ATAC-seq sample preparation

Nuclear isolation, tagmentation and PCR amplification was carried out according to Buenrostro *et al.* [28]. In brief, 50000 cells per sample were transferred to 0.2 ml tubes, centrifuged (500xg, 6 min, 4°C) and the supernatant was removed. Cells were washed with PBS (500xg, 6 min, 4°C) and lysed in lysis buffer (10 mM Tris-HCl, pH 7.4; 10 mM NaCl; 3 mM MgCl_2_; 0.1% IGEPAL CA-630) to isolate nuclei. 3 technical replicates (50000 cells each) were processed in parallel for each sample (except donor A, 2 technical replicates). Nuclei were washed in PBS (500xg, 6 min, 4°C), and resuspended in Transposase Reaction mix. Transposition was carried out at 37 °C for 30 minutes, followed by clean up using Qiagen Minelute Reaction Clean Up kit according to the manufacturer’s instructions (Qiagen). PCR amplification using reagents from Nextera DNA Sample Preparation Kit (Illumina) and barcoding of replicates was performed with reaction conditions and index primers as described in [28]. 20 different index primers were used and distributed across replicate samples and donors in a balanced way to control for potential batch effects. The PCR product was cleaned up using Qiagen Minelute Reaction Clean Up kit. Subsequently, gel size selection was performed by gel electrophoresis [1.8% (w/v) certified low-melt agarose (Bio-Rad) in 1x UltraPure Tris Acetate-EDTA (TAE) buffer (Invitrogen, Thermo Fisher Scientific), with a free well between each of the samples] and DNA in the size range of 150-230 bp was excised using surgical blades. Replicate samples were allocated to the gels in a balanced way regarding donor and experimental condition to control for potential batch effects. Resulting DNA libraries were purified using Qiagen Minelute Gel Extraction Kit according to the manufacturer’s instructions (Qiagen).

Size distribution of ATAC-seq sequencing libraries was determined on a 2100 Bioanalyzer instrument (Agilent Technologies) using the Agilent DNA High Sensitivity Kit according to the manufacturer’s instructions. Libraries were quantified by real-time PCR on a StepOne plus detector system (Applied Biosystems) using the KAPA Library Quantification Kit (KAPA Biosystems). Sequencing was performed on an Illumina HiSeq 2500 instrument (Illumina) with single-read setting and read length 42 bp.

#### RNA-seq sample preparation

Cells were centrifuged (1000xg, 5 min, 20°C), washed with PBS, and lysed in QIAzol Lysis Reagent (Qiagen) by vortexing and incubating for 5 min at 20°C. Lysates were stored at −80°C until RNA extraction. RNA was extracted with the miRNeasy Micro Kit (Qiagen) according to the manufacturer’s instructions. RNA concentration was determined on a Nanodrop 2000 spectrophotometer (Thermo Fisher Scientific) and RNA quality was controlled on a 2100 Bioanalyzer instrument (Agilent Technologies) using an Agilent RNA 6000 Pico Kit. RNA integrity numbers (RIN) were 8.5±0.3 (mean±SD). Libraries were prepared in one batch with the TruSeq Stranded mRNA HT kit (Illumina) with Dual Indexing adapters and Ambion ERCC Spike-In Control (Thermo Fisher Scientific). Samples were allocated to sequencing indexes and -lanes in a balanced fashion to control for potential batch effects (7-8 samples/lane). Library concentration was determined on a Qubit 2.0 Fluorometer (Thermo Fisher Scientific). Library size and quality were measured on a 2100 Bioanalyzer instrument (Agilent Technologies) using an Agilent High Sensitivity DNA Kit. Libraries were quantified with the KAPA Library Quantification Kit (KAPA Biosystems). Sequencing was performed on a HiSeq 2500 Sequencing Platform (Illumina; High Output Run) with 76 nt paired-end reads.

### Computational Methods

#### Preprocessing of sequencing data

BCL base-call files were demultiplexed and converted to FASTQ files using bcl2fastq version 2.17.1.14. For quality control FastQC version 0.11.5 was used.

FASTQ files from the ATAC-seq experiments were trimmed from adapters and low quality bases using scythe version 0.991. and sickle version 1.33. FASTQ files from the ATAC-seq experiments were aligned to the human genome version hg38 using bowtie version 2.3.0 with the ‘--very-sensitive’ option. After alignment, BAM files of ATAC-seq experiments were filtered to eliminate: duplicates (samtools rmdup), alignments with a mapping score below 10, and alignments that are not mapped to chromosome 1-22, chromosome X or Y. ATAC-seq peaks were called using the ‘findPeaks’ script from the HOMER suite (version 4.9.1) with the ‘-style factor’ option [41]. ATAC-seq peaks were assigned to a gene using the ‘annotatePeaks’ script from the HOMER suite.

FASTQ files from the RNA-seq experiments were aligned using STAR version 2.5.2b. Indexes for RNA-seq alignment were created using the gencode version 25 annotation file. RNA-seq alignment was run with STAR’s built-in adapter trimming option (‘--clip3pAdapterSeq AGATCGGAAGAGCACACGTCTGAACTCCAGTCAC AGATCGGAAGAGCGTCGTGTAGGGAAAGAGTGTA’) and its built-in counting option (‘-- quantMode’). Only genes with more than 1 count per million in at least 3 samples were included in the downstream RNA-seq analysis.

#### Generation of consensus peak-set

In order to generate a set of comparable features (genomic regions) for read-counting and quantifying differential accessibility from the ATAC-seq data, a set of consensus peaks was generated in two subsequent steps. (1) Generation of consensus peaks on the technical replicate level: First, the peaks that appeared in at least two technical replicates (out of a total of three, except one donor with a total of two technical replicates) with at least 75% reciprocal overlap were selected. Then these selected regions were partitioned into disjoint non-empty subsets so that each element is contained in precisely one subset. Only the partitions appearing in at least two replicates were retained, and afterwards adjacent regions were merged. A bed file resulting from these steps is herein referred to as a sample. (2) Next, all bed files containing each set of technical replicate level consensus regions were concatenated and the presence of each region was counted within each experimental sample; one occurrence corresponds to one donor (biological replicate) and one cell type (experimental condition). Only the regions appearing in at least four samples were kept (n=5 is the number of biological replicates in the smallest group regarding experimental condition). Regions having a distance of 42 bases or less between them were subsequently merged (the number 42 corresponds to the sequencing read length in bases). Afterwards, reads were counted using the featureCounts tool [42] with the criterion that at least half of the read had to overlap with a feature to be assigned.

#### Calculation of differential expression and differential accessibility

Differential expression and accessibility was calculated using the edgeR [43] library version 3.18.1 from Bioconductor. Donors (biological replicates) and cell types (experimental conditions) were used as explanatory variables in the generalized linear models. ATAC-seq data were normalized to length and GC content by conditional quantile normalization (CQN) [44]. Comparisons were made between GM-CSF positive CD4 T cells *versus* GM-CSF negative CD4 T cells, or between memory *versus* naïve CD4 T cells (see Figure 1A). The cutoff to call differentially expressed genes (DEGs) or differentially accessible regions (DARs) was FDR<0.05 and >25% fold change (in the direction of either up- or down-regulation that is, either 1.25 or 0.75 fold change).

#### Footprinting

Footprinting was carried out using the Wellington algorithm [45], i.e. the wellington_footprints.py script from the pyDNase library version 0.2.5 with the following settings: -fp 6,41,1 -sh 7,36,1 -fdr 0.01 -fdriter 100 -fdrlimit −30 -A. The footprint occupancy score (FOS) for each footprint was calculated using the pyDNase library as described in [46]. For subsequent network reconstruction, we considered only footprints with a FOS smaller than the following threshold: median(FOS) + [median(FOS) - minimum(FOS)].

#### Network reconstruction

A directed network was reconstructed by combining information from the ATAC-seq and RNA-seq data in two subsequent steps. (1) Identifying source nodes: Peaks were ranked based on their combined measure of significance and direction of differential accessibility [− log_10_(FDR) × sign(log_2_(fold change))]. Peaks containing footprints were scanned for TF binding motifs using the TRANSFAC database [47]. An enrichment score was calculated to identify TFs with binding sites enriched in differentially accessible peaks, using tools similar to gene set enrichment analysis (GSEA) [48]. In detail, random sampling was performed on the ranked list of peaks to assess how probable it is to observe at least the same enrichment by chance (P value) [49]. After multiple test correction, TFs with FDR<0.05 were selected and the normalized enrichment score (NES) was obtained. Only those source nodes (TFs) that were detectably expressed on RNA level (according to a minimal RNA-seq filtering rule) were considered. Of these, most fell into the class of highly expressed genes (HEGs) according to [50]. (2) Identifying target nodes: Target nodes are defined as peaks with an assigned gene. The selection criteria were that (i) the peak contains a footprint with a binding motif of the source node (TF) and (ii) the peak and/or the assigned gene has to be differentially accessible or differentially expressed respectively.

To assign an importance measure to the source nodes in networks generated as above, the PageRank [51] of the network nodes was calculated after inverting the directionality of all edges in the network (only for the purpose of this computation). After this computation, the nodes with high PageRank values (higher than the 99th percentile of all node values within a given network) were selected from both the GM-CSF and memory network, and afterwards their values were investigated in each of the two networks.

## RESULTS AND DISCUSSION

GM-CSF positive CD4 T cells are enriched in MS patients and play a crucial role in EAE; nevertheless the factors driving and markers defining those cells are largely unknown. To better understand the features and regulatory networks of GM-CSF positive CD4 T cells, we therefore studied the transcriptional profiles and chromatin accessibility of these cells. *In vitro* differentiated GM-CSF producing cells comprise several subsets [27] and are likely to differ from those generated *in vivo*. Further, *ex vivo* re-stimulation with strong artificial stimuli such as PMA and ionomycin - which is usually necessary to reach sufficient signal strength for detection by intracellular cytokine staining - drastically alters the transcriptome of T cells. Hence, we aimed to isolate GM-CSF producing cells *ex vivo* in an as much as possible unmanipulated state by GM-CSF secretion assay, “capturing” and isolating those cells that actively secrete GM-CSF (experimental setup, see Figure 1A), here defined as GM-CSF positive cells. The capture assay was performed starting from highly purified CD4 T cells derived from human peripheral blood (purity 97.3±0.6 %, mean±SEM, Supplementary Figure S1A). As controls, we used the respective GM-CSF negative fraction from the isolation procedure, as well as the bulk CD4 T cells before any capture assay procedure. The latter should, given the low fraction of GM-CSF positive cells, be very similar to the GM-CSF negative fraction and hence allows for estimation of the effects arising from the capture assay procedure. The purity of GM-CSF positive and GM-CSF negative fractions was assessed by flow cytometry (Figure 1B) and the yield of isolated GM-CSF positive cells was 1.2±0.6 % (mean±SD) of CD4 T cells. To measure the transcriptome and DNA accessibility from limited cell numbers, we employed highly sensitive next-generation sequencing (NGS) methods (RNA-seq and ATAC-seq respectively). Since cytokine secreting cells may differ from naïve cells due to a memory-like phenotype, and a large fraction of CD4 T cells are naïve (mean±SEM 42.3±5.8 % in the donors here; see Supplementary Figure S1A), we further profiled highly purified naïve and memory CD4 T cells from the same donors (Figure 1A and Supplementary Figure S1B-D).

Altogether, we obtained DNA accessibility and transcriptome data from highly purified *ex vivo* derived human naïve CD4 T cells, memory CD4 T cells, bulk CD4 T cells and well as GM-CSF positive and corresponding GM-CSF negative CD4 T cells. To enable paired analysis within a donor, these cell populations were isolated in parallel within a donor; for 6 donors in total. DNA accessibility and mRNA data were obtained in parallel from the same samples allowing for matched integration of the data.

### Unique and shared DNA accessibility and gene expression signatures of GM-CSF positive and memory CD4 T cells

We studied the above-described five different CD4 T cell populations by RNA-seq and ATAC-seq. To minimize potential batch effects due to technical factors, the library preparations and sequencing runs were designed in such a way that donors, cell populations, and (for ATAC-seq) technical triplicates were distributed in a balanced fashion. It is also worth noting that only donors of the same gender were studied here (male, age 35.7±8.5 years, mean±SD), which may be important since it was recently shown that gender was the largest source of variation explaining chromatin accessibility in primary human CD4 T cells measured by ATAC-seq [36]. That study further discovered novel elements escaping X chromosome inactivation and affecting immune genes [36]. To assess which factors explained most of the variability between the samples under study here, we performed principle component analysis (PCA). Indeed, for both data types there was a grouping of the samples based on the cell subset, outweighing donor or experimental variation (Figure 2A, B) and confirming the quality of our samples and data. Notably, for RNA data, the cell populations were generally more distinct from each other than for DNA accessibility data. However, the GM-CSF positive and corresponding GM-CSF negative fraction appeared relatively similar to each other in the PCA performed on RNA data, while PCA results from ATAC-seq data were closer to the expected pattern that is, bulk CD4 T cells appearing “between” GM-CSF positive and GM-CSF negative populations (Figure 2A, B). The difference between RNA-seq and ATAC-seq data with respect to separation of GM-CSF positive and negative cells may indicate that the capture assay procedure imposes distinct changes on the transcriptome, highlighting the importance of using correspondingly treated controls to determine differential expression. In contrast, changes in DNA accessibility appeared more robust towards changes due to the experimental procedure at least within the experimental time frame under study, although the distinction of the other groups was generally less apparent with ATAC-seq data. Because the first two PCs only explained about 50% of the variation in the data, we also used another dimensionality reduction method to explore the sample-to-sample relationships, namely t-distributed stochastic neighbor embedding (t-SNE) [52]. t-SNE, in contrast to PCA, is a non-linear dimensionality reduction algorithm, and it is suited for capturing local and global relationships at the same time. The t-SNE results (Figure 2C, D) generally confirmed the results of the PCA analysis (Figure 2A, B) that is, the groups (cell types) being more distinct in RNA data than ATAC-seq data, with the exception of GM-CSF positive and GM-CSF negative cells.

**Figure 2.**
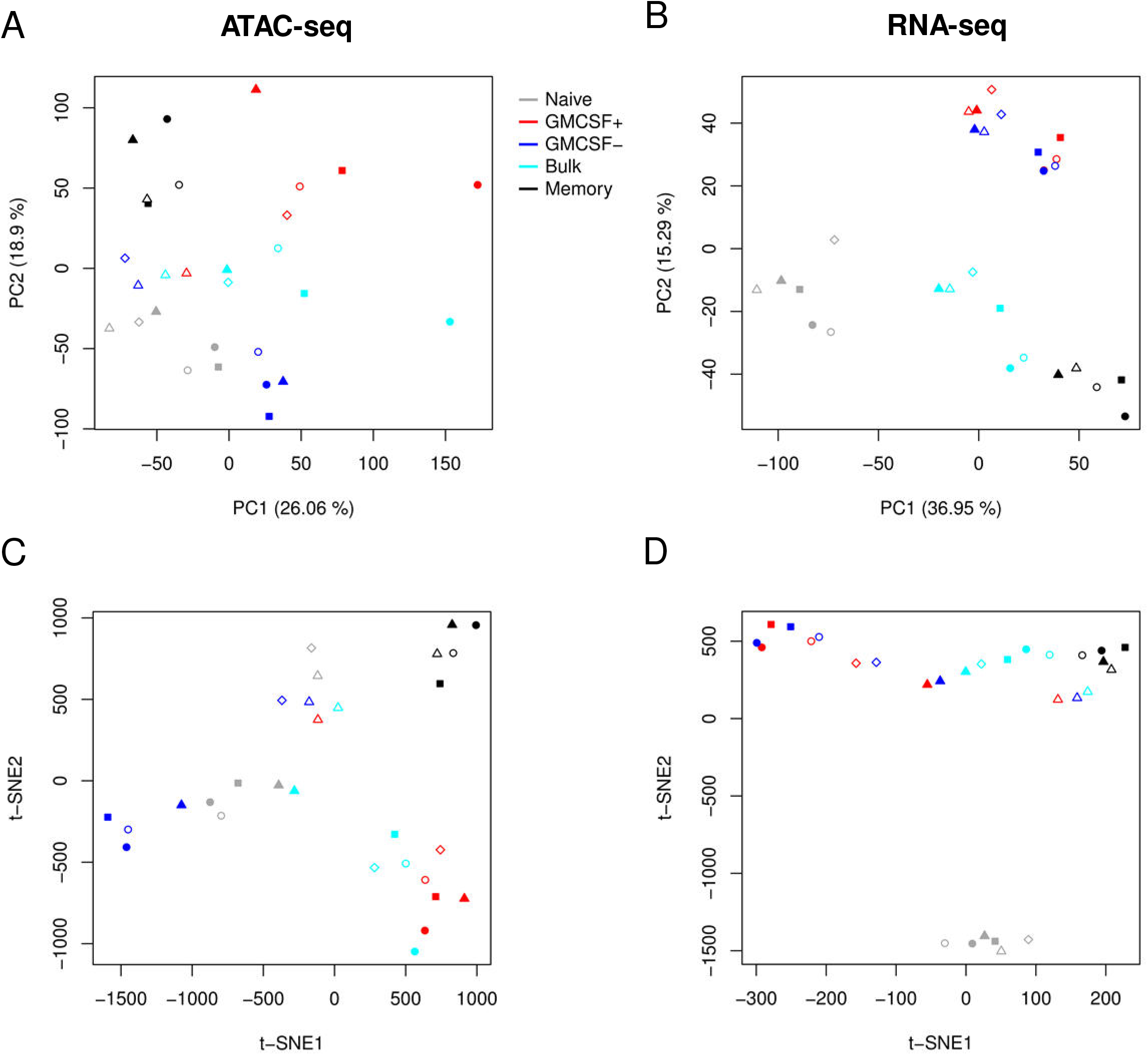
Explorative analysis of ATAC-seq and RNA-seq samples. (**A**) Principal Component Analysis (PCA) was carried out on CQN-normalized ATAC-seq data given as log_2_(RPKM+1) centered by mean subtraction for each feature (genomic region) across samples. PC1 and PC2 are shown, along with the % variation explained. Symbol colors indicate the given cell populations, and symbol shapes and fillings represent individual donors (n=5-6 donors) as in Figure 1B. Data from technical replicates were pooled before analysis. (**B**) PCA for RNA-seq data given as log_2_(FPKM+1) centered by mean subtraction for each feature (gene) across samples. Labels as in (A). (**C, D**) t-SNE dimensionality reduction visualization of ATAC-seq (C) and RNA-seq (D) data, processed and labeled as in (A) and (B), respectively.

We defined significantly differentially accessible DNA regions (DARs) and significantly differentially expressed genes (DEGs) in GM-CSF positive CD4 T cells or in memory CD4 T cells. To do so, we specifically analyzed the signatures of GM-CSF positive *versus* GM-CSF negative CD4 T cells, as well as the profiles of memory *versus* naïve CD4 T cells (Figure 1A and Figure 3A, B). We used generalized linear models based on the negative binomial distribution (edgeR) to determine differential expression and accessibility respectively, and we called DEGs and DARs based on combined FDR and fold change cutoffs. We called 16571 DARs in GM-CSF positive CD4 T cells (compared to corresponding GM-CSF negative cells; Figure 3A) and 13705 DARs in memory CD4 T cells (compared to naïve CD4 T cells; Figure 3A). On the transcriptome level, we called 124 DEGs in GM-CSF positive and 5383 DEGs in memory CD4 T cells (Figure 3B). The relatively low number of DEGs in GM-CSF positive cells is in agreement with the PCA and t-SNE data (Figure 2B, D) and may suggest that combination with ATAC-seq data drastically improves the possibility to define molecular signatures specific to GM-CSF positive *ex vivo* captured cells.

**Figure 3.**
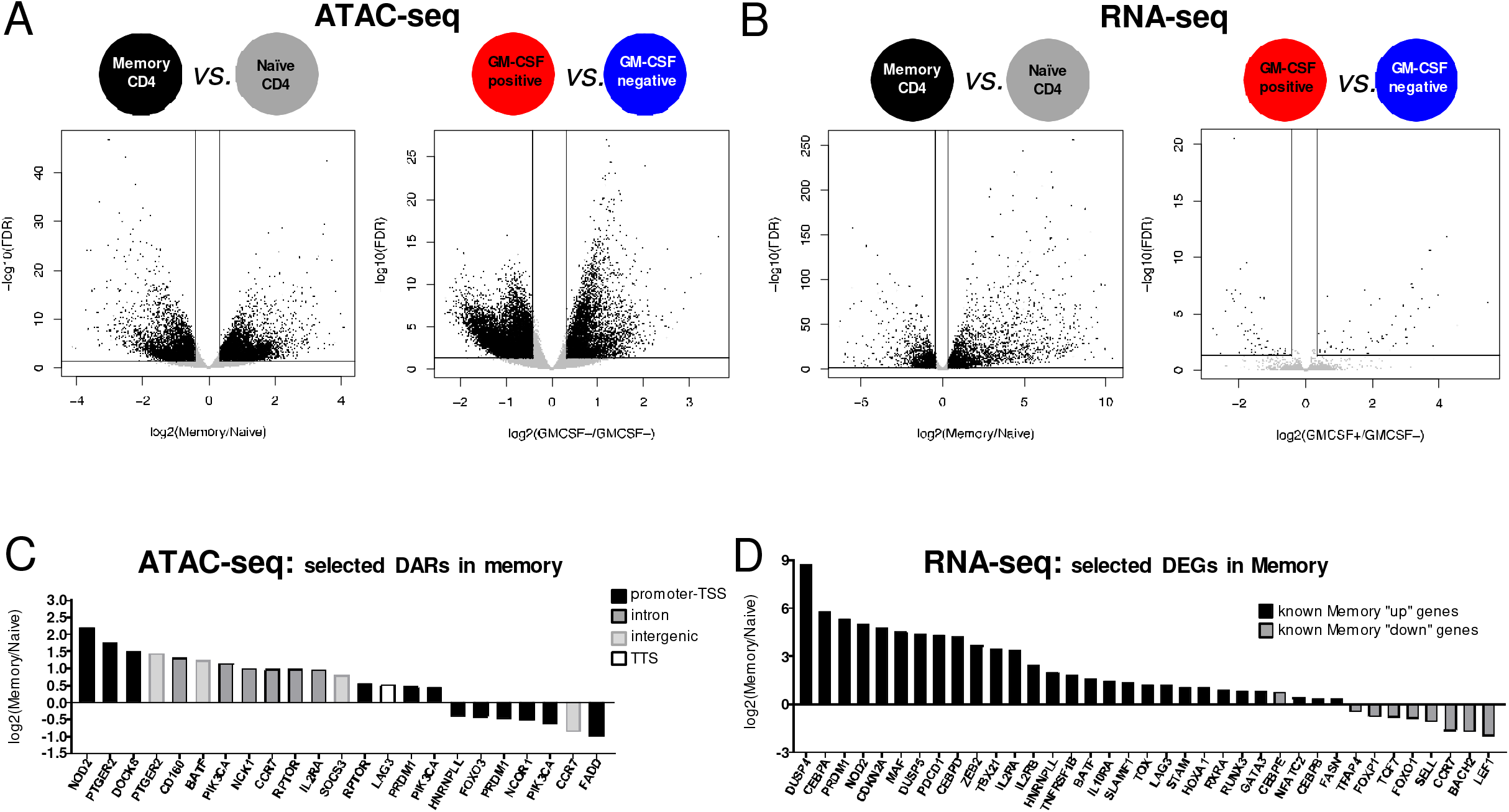
Differential chromatin accessibility and gene expression in memory *versus* naïve CD4 T cells as well as in GM-CSF positive *versus* negative cells. (**A**) ATAC-seq data were CQN-normalized and differential accessibility between the indicated cell population comparison was calculated (left: memory *vs.* naïve CD4 T cells; right: GM-CSF positive *vs.* negative CD4 T cells). Volcano plots show each consensus peak as a single dot, and lines indicate the threshold for calling a DAR (FDR<0.05 and >25% fold change). Not significantly differentially accessible regions are depicted in grey. (**B**) Differential gene expression was calculated from RNA-seq data for cell populations as in (A). Lines indicate the threshold for calling a DEG (FDR<0.05 and > 25% fold change), not significantly differentially expressed genes are depicted in grey. (**C**) A selection of DARs in Memory *vs.* Naïve (not DAR in GM-CSF positive *vs.* GM-CSF negative) was studied for being assigned to genes known to play a role in T cell memory. Log_2_(fold change(Memory/Naïve)) for selected DARs (FDR<0.05) are plotted, and colors indicate the category the respective region is assigned to (TSS: transcription start site; TTS: transcription termination site). (**D**) Known T cell memory “up” (black) or T cell memory “down” (grey) genes based on previous literature were selected if differentially expressed (FDR<0.05 in Memory *vs.* Naïve) in the present data. log_2_(fold change(Memory/Naïve)) for these selected DEGs is plotted; values >0 represent up-, values <0 represent down-regulation in Memory T cells. Colors indicate whether the gene was previously described to be “up” or “down” in memory T cells.

Next, we studied the DARs and DEGs in more detail. We first focused on the signatures of memory CD4 T cells, which are well studied in the literature [29] and hence enabled to assess the biological quality of our data, besides providing a new NGS data set of human primary memory and naïve CD4 T cells. DARs and DEGs defined in memory cells are shown in Supplementary Figure S2, along with their molecular patterns in the other cell types under study. We next extracted those memory-specific DARs that were assigned to a gene from a list of genes known to be involved in T cell memory as compiled by Polansky and colleagues [30]. Several of the DARs in memory cells were assigned to such known memory-associated genes, most of those in the promoter region (Figure 3C). Furthermore, we assessed a selected subset of memory related genes that were shown to be up- or down-regulated on RNA level in memory T cells [30]. The majority of these genes were DEG in memory cells in our data, notably up- or down-regulated almost exclusively (36 of 37 studied genes; 97%) in the expected direction (Figure 3D), validating our data.

### The molecular signature of GM-CSF positive CD4 T cells

Next, we focused on the signatures of GM-CSF positive cells by studying the respective DARs and DEGs in more detail. All the DARs in GM-CSF positive cells are displayed in Figure 4A with the color scale representing accessibility. A substantial fraction of DARs in GM-CSF positive cells displayed a similar pattern in memory cells, while naïve cells and GM-CSF negative cells were most distinct from GM-CSF positive cells (Figure 4A). In bulk CD4 T cells, DARs defined in GM-CSF positive cells showed heterogeneity between donors (Figure 4A), which may reflect the variability in the fraction of memory and naïve cells within bulk CD4 T cells depending on the donor (Supplementary Figure S1A). Importantly, several DARs displayed a unique accessibility pattern in GM-CSF positive cells distinct from other cell populations under study (Figure 4A). These data show that GM-CSF positive cells can be assigned a specific pattern of accessible DNA regions that distinguish them from other CD4 T cell subsets, and that may contribute important information about regulation of GM-CSF positive cells. Different DNA accessibility can functionally affect the status of a cell by, for example, modifying expression of genes regulated by these regions.

**Figure 4.**
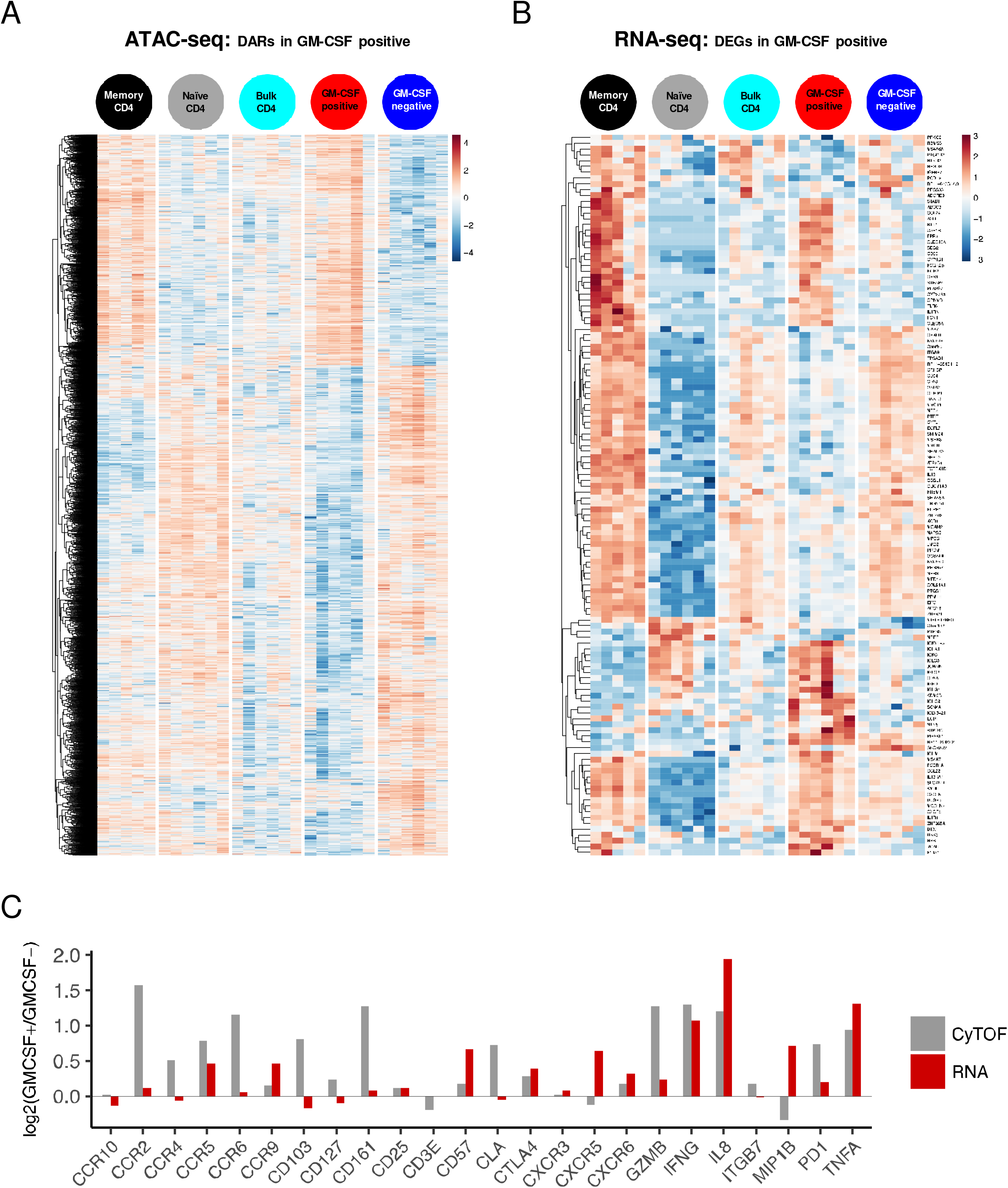
Expression profile and DNA accessibility signatures of human GM-CSF positive CD4 T cells. (**A**) Differentially accessible DNA regions in GM-CSF positive *versus* GM-CSF negative CD4 T cells (FDR<0.05 in this comparison) are plotted as a heatmap for their accessibility in all the five cell populations studied. CQN normalized log_2_(RPKM+1) is represented by the color scale, indicating accessibility (blue: low, red: high). Data were row-scaled and -clustered (Euclidean distance, complete linkage clustering). (**B**) Differentially expressed genes (RNA) in GM-CSF positive *versus* GM-CSF negative CD4 T cells (FDR<0.05 in this comparison) are plotted as a heatmap for their expression in all the five cell populations studied. Gene expression is displayed as log_2_(FPKM+1) with blue indicating low and red indicating high expression according to the color scale. Data were row-scaled and -clustered (Euclidean distance, complete linkage clustering). (**C**) The mean of log_2_(fold change) of GM-CSF+/GM-CSF- cell populations using the median intensity values from CyTOF measurements are shown, using CD4 T cell-gated PBMC data from Wong *et al.* [57] and designated as “CyTOF” (grey bars). For the genes corresponding to the proteins measured in CyTOF, the log_2_(fold change) of GM-CSF+/GM-CSF- cell populations from the RNA-seq data of this study is plotted, calculated with the generalized linear model (edgeR) and labelled as “RNA” (red).

Thus, we then studied the DEGs defined in GM-CSF positive cells by analyzing their expression in GM-CSF positive cells along with the other CD4 T cell subsets under study. Like observed for the DARs, DEGs in GM-CSF positive cells shared a large part of the RNA signature with memory cells, but also displayed distinct patterns, and differed largely from naïve cells, GM-CSF negative and bulk CD4 T cells (Figure 4B). Despite the general similarity of cells treated with the capture assay regarding the global transcriptome (Figure 2B, D), sub-setting on the DEGs defined between GM-CSF positive and negative cells with stringent statistical cutoffs visualized clearly the differences in these “signature genes” for those cell types (Figure 4B), albeit this is not surprising due to the respective pre-selection of the genes. However and importantly, sub-setting on these genes that were defined as DEGs of GM-CSF positive *versus* corresponding negative cells (that is, without considering the bulk CD4 T cell samples) also clearly showed the expected similarity of bulk CD4 T cells and GM-CSF negative CD4 T cells (Figure 4B) that was not apparent in the RNA-seq PCA on all genes (see above; Figure 2B, D), confirming that the selected GM-CSF positive cell “signature” DEGs are likely not affected by the capture assay and column procedure. DEGs in GM-CSF positive cells comprised multiple genes with a well-known role in T cells such as *EGR2, IFNG, CTLA4, CXCL8* and *CXCR5* along with multiple genes with an unknown role in T cells and potentially regulating GM-CSF positive cells (Figure 4B). *CSF2RB* (encoding for the high affinity receptor subunit for IL-3, IL-5 and GM-CSF) was among the DEGs in GM-CSF positive cells, suggesting a possible feedback loop; *CSF2RB* expression being lower in GM-CSF positive than negative cells excludes a technical artifact of isolation of cells binding GM-CSF through this receptor (Figure 4B). Albeit not passing the significance threshold for being called a DEG in GM-CSF+ cells, relatively high expression of *IFNG* in GM-CSF positive cells (data not displayed) is in accordance with our and other’s previous findings from flow cytometry of T cells from healthy donors as well as MS patients demonstrating preferential co-expression of IFN-γ and GM-CSF in some, but not all GM-CSF producing human CD4 single T cells or clonal populations thereof [11,14,26,27,53]. Also in line with the majority of these studies of human T cells, *IL17A* and *IL17F* expression was below detection limit in GM-CSF positive cells (as well as any other cell population under study; data not displayed). It should be noted that *CSF2* mRNA, which encodes for GM-CSF, was lowly expressed in all samples (RNA-seq counts close to detection limit), which may be explained by rapid mRNA decay conferred by the adenine and uridine-rich elements (ARE) in the GM-CSF promoter – AREs in fact have been discovered in the *CSF2* gene which codes for a particularly unstable transcript [54–56]. Importantly, considering the instability of *CSF2* mRNA and relatively low expression levels, our approach of isolating GM-CSF protein secreting cells is likely to be more suitable to define signatures of *ex vivo* derived GM-CSF positive cells, as opposed to for example single cell RNA-seq of mixed T cell populations.

Since to our knowledge, there is no other data set available that studied the signatures of purified GM-CSF secreting cells, the RNA expression signatures defined in the GM-CSF positive cells in this study could not be validated externally in an independently published transcriptome data set. Nevertheless, we strived to confirm the expression signature from GM-CSF positive captured cells by comparing with a completely independent data set and experimental setup. To this end, we asked whether the RNA expression pattern of captured GM-CSF positive T cells generally agreed with the respective proteins in GM-CSF positive T cells defined by intracellular cytokine staining. To explore overlap of as many as possible markers, we studied a mass cytometry (CyTOF) data set [57] which comprises staining of human PBMCs with CD4 and GM-CSF along with other markers measured on protein level. In pre-gated CD4 positive T cells of this CyTOF data set, we gated on GM-CSF+ cells and GM-CSF- cells and determined the relative expression of other available protein markers in these populations. We compared the up- or down-regulation of these protein markers in gated GM-CSF+ *versus* GM-CSF- cells [log_2_(fold change) of the median signal intensity of the two cell populations] with the up- or down-regulation of the corresponding protein-coding transcripts of these markers in isolated GM-CSF positive *versus* GM-CSF negative cells from our data [log_2_(fold change) of mRNA expression]. As a result, 16 of the 24 markers (two thirds) had a fold change with identical directions in the CyTOF and RNA-seq data (Figure 4C). According to a binomial distribution, the probability of observing this or greater concordance between the two datasets by chance is 7.6%. This needs to be acknowledged considering that those markers that do not concur might be regulated by protein internalization from the surface, such as well-known for CD3 [58] that was down-regulated in CyTOF data but barely affected on RNA level. Also, the total abundance of certain proteins may be regulated on post-transcriptional level, as transcript levels cannot always predict protein abundance [59,60]. Therefore, we concluded that the expression profile of the GM-CSF positive captured T cells matches well with the profile of independently characterized human GM-CSF positive T cells.

### GM-CSF positive CD4 T cell transcript signatures and chromatin accessibility are associated with autoimmune diseases, especially MS

Next, we asked whether the gene expression pattern of GM-CSF positive cells was enriched for any diseases by exploring the Open Targets platform [61]. We ranked the detected genes based on their differential expression in GM-CSF positive *versus* negative cells using the −log_10_(FDR) × sign(log_2_(fold change)) function for ranking, and calculated enrichment scores for diseases. Indeed, among the few significantly enriched diseases (FDR<0.05), the autoimmune diseases MS and rheumatoid arthritis were represented (Figure 5A). Notably, for both diseases GM-CSF targeting is in clinical trials [15], supporting the relevance of our data. The data also showed enrichment for several other diseases related to the immune system, infection or metabolism, although it should be noted that some of these contained only few elements (genes) and may thus be less relevant than MS or rheumatoid arthritis, which comprised a large number of genes (Figure 5A and Supplementary Table S1A). When performing the same enrichment analysis for the ranked gene list from memory *versus* naïve CD4 T cells, a large number of diseases were significantly enriched including many diseases involving the immune system, as expected (Supplementary Table S1B).

**Figure 5.**
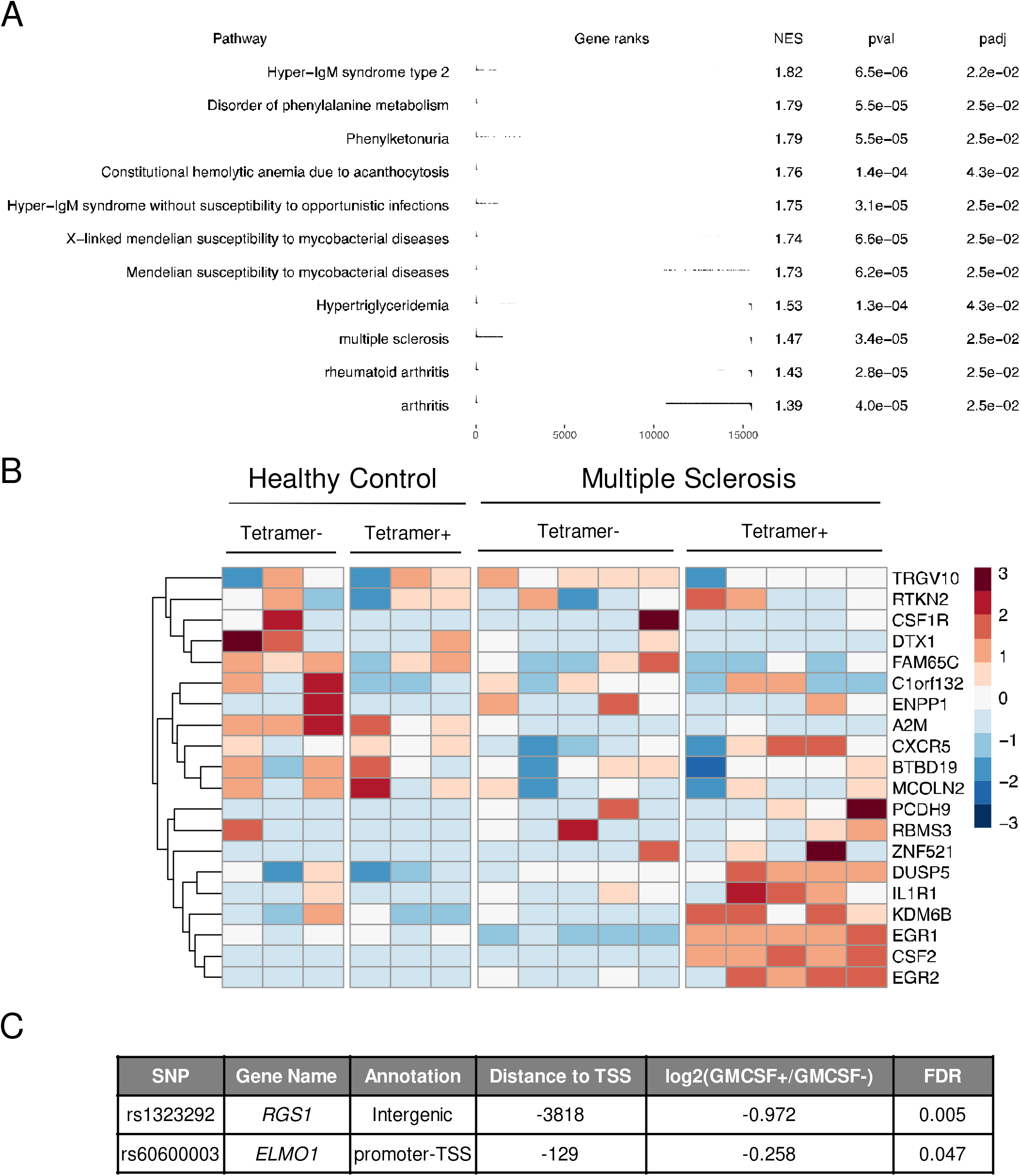
GM-CSF positive CD4 T cell signatures are associated to autoimmune diseases, especially MS. (**A**) Rank based gene set enrichment analysis is shown using the −log_10_(FDR) × sign(log_2_(GM-CSF+/GM-CSF-)) function to rank genes based on differential expression according to the GM-CSF+/− cell populations, and the Open Targets database [61] was used to provide lists of genes associated with diseases. NES: normalized enrichment score; pval: P value; padj: FDR. (**B**) The Heatmap represents the row-scaled log_2_(RPKM+1) expression values from RNA-seq data of CD4 T cells from MS patients or healthy controls (data from [26]). Groups are separated based on disease status (MS or healthy) and myelin antigen reactivity (reactive: Tetramer+). Genes are selected as those identified in the current RNA-seq study as differentially expressed between GM-CSF positive *versus* GM-CSF negative cells and having detectable expression (log_2_RPKM>0) in at least 4 samples in the data from [26]. (**C**) Of the 110 established non-MHC MS susceptibility variants [63], two SNPs mapped to a consensus peak from the current ATAC-seq study. Information about these SNPs is shown in the table, along with the differential accessibility analysis in GM-CSF positive *versus* GM-CSF negative CD4 T cells.

Due to the relevance of GM-CSF positive T cells in MS, we studied in more detail whether DEGs identified in GM-CSF positive cells displayed altered expression in MS patients’ T cell samples. T cells recognizing peptides derived from myelin proteins as autoantigen and being activated and migrating to the CNS are thought to be crucial mediators of inflammation in MS [62]. Hafler and colleagues have performed RNA-seq of expanded CD4 T cells derived from MS patients and healthy controls, with two subsets of samples each containing those auto-reactive to antigenic peptides derived from myelin (tetramer positive) *versus* tetramer negative T cells [26]. The authors discovered that myelin reactive T cells from patients with MS displayed strongly enhanced production of IFN-γ, IL-17 and GM-CSF compared to those isolated from healthy controls, which instead secreted more anti-inflammatory IL-10 [26]. We studied whether these cells would also display altered expression of the genes we defined as signature genes of GM-CSF positive cells. Indeed, a subset of these genes was down- and another subset was up-regulated in myelin-reactive T cells from MS patients (Figure 5B), potentially identifying genes relevant to the disease pathogenesis. Here, the T cells displayed detectable *CSF2* mRNA encoding for GM-CSF, perhaps due to the *in vitro* stimulation and expansion of these T cells. Remarkably, only myelin reactive cells from MS patients expressed high levels of *CSF2* (Figure 5B). Interestingly, this *CSF2* expression pattern strongly resembled the expression patterns of several of the GM-CSF positive T cell signature DEGs defined here, namely *DUSP5*, *IL1R1*, *KDM6B*, *EGR1* and *EGR2* (Figure 5B) which may be interesting candidates to explore in the future regarding their role in GM-CSF positive T cells and MS. While this analysis confirmed the relevance of the GM-CSF positive T cells’ RNA-seq data in MS, we next analyzed whether also the ATAC-seq data could reveal information on those DNA regions most relevant for GM-CSF positive T cells in the context of MS. To do so, we asked whether the peaks called from our ATAC-seq analysis contained any SNPs associated with MS. Interrogating the current best list of non-MHC MS susceptibility variants comprising 110 established risk variants from the International Multiple Sclerosis Genetics Consortium [63,64], we identified two SNPs mapping to the accessible DNA peaks (considering all consensus peaks) from our study. These SNPs were assigned to the protein-coding genes *Regulator Of G Protein Signaling 1* (*RGS1*) and *Engulfment And Cell Motility 1* (*ELMO1*) genes respectively (Figure 5C). Importantly, the regions containing these SNPs were significantly differentially accessible (FDR<0.05) in GM-CSF positive *versus* GM-CSF negative cells (Figure 5C), suggesting a putative role of these regions in T cell mediated MS pathogenesis.

### Relationship of differential gene expression and chromatin accessibility

Since chromatin accessibility can directly affect gene expression, we next combined the ATAC-seq and RNA-seq data, aiming to identify key TFs that may bind to open chromatin regions and hence affect the expression of their target genes. The methods for calling consensus peaks are not trivial. Therefore, we first confirmed that the genes assigned to the consensus peaks defined in this study were enriched for several pathways involved in immune regulation including T cell receptor signaling and Th subset differentiation, as well as for immune-related diseases including MS, RA and other autoimmune disease (Supplementary Table S2A, B). Furthermore, the distribution of the consensus peaks (open chromatin regions) defined in our data showed an enrichment for being located in CpG islands, promoters, 5’ UTRs, exons and protein-coding regions (Figure 6A), suggesting that the ATAC-seq data generated here should be well suited to identify TF binding and the expression of corresponding target genes. We first studied the relationship of RNA and chromatin data on a global level: We considered the genes that were both detected on RNA level and also had an ATAC-seq peak assigned to them. We ranked these genes using the −log_10_(FDR) × sign(log(fold change)) function separately for the RNA-seq and ATAC-seq data. Next, we visualized the correlation between these two ranks for each contrast (Figure 6B). Considering only the genes significantly changing (FDR<0.05) in the given contrast and assigning them to up- and down-regulated categories, we detected more than random coincidence in the direction of the change between the RNA-seq and ATAC-seq data (using Fisher’s exact test with Monte Carlo simulation; Figure 6B). Increased openness of the chromatin was associated with increased expression of the corresponding gene’s RNA for the majority of genes. Less accessible chromatin also coincided with low expression of the corresponding gene in many cases, although a substantial fraction of lowly accessible regions also displayed high expression of the corresponding gene (Figure 6C).

**Figure 6.**
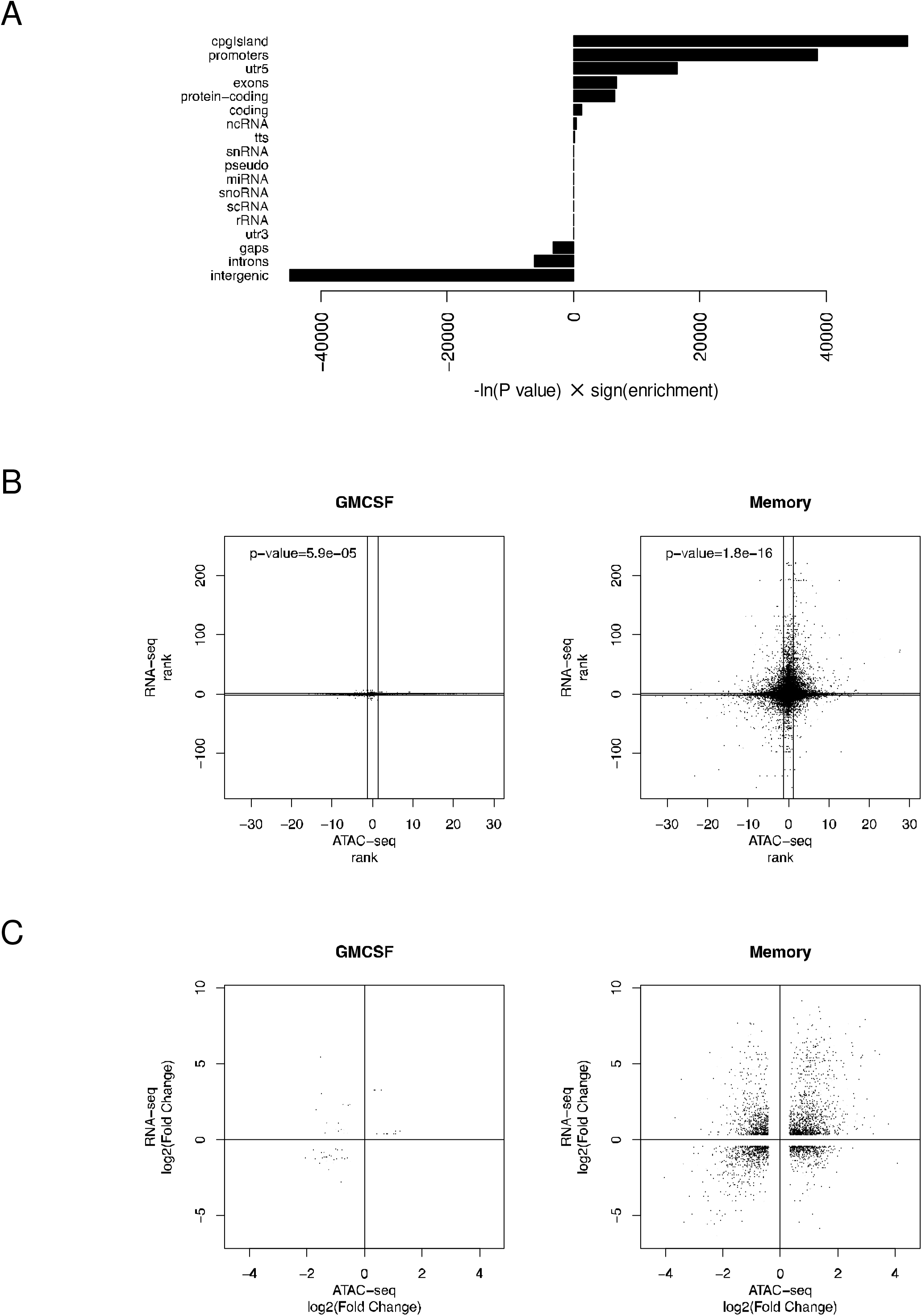
Enrichment of ATAC-seq peaks in regions representing certain genomic features, and relationship of ATAC-seq and RNA-seq data. (**A**) The enrichment of ATAC-seq consensus peaks in different types of genomic regions is shown with bars representing the enrichment significance × direction of enrichment (-ln(P value) × sign(enrichment)). (**B**) Genes that were detected on both RNA-seq level and were also assigned to ATAC-seq peaks, were ranked using the −log_10_(FDR) × sign(log_2_(fold change)) function based on both RNA-seq and ATAC-seq (using values of the assigned peaks) within a given cell type comparison (left: GM-CSF+ *vs.* GM-CSF-; right: memory *vs.* naïve). The ranks of genes using ATAC-seq *vs.* RNA-seq for ranking are visualized for both cell type comparisons; lines indicate FDR cutoff 0.05 in the respective comparison. P value represents the probability of observing this or more directional agreement between the two datatypes by chance, using only genes significantly differential (FDR<0.05) in both data types. P values were calculated by Fisher’s exact test with Monte Carlo simulation. (**C**) Dot plot representation as in (B), but using log_2_(fold change) for ranking, and visualizing only (assigned) genes that are significantly differential in both data types.

### Identification of key TFs linked to the signatures of GM-CSF positive and memory CD4 T cells

To identify potential TFs that may establish the gene signatures of GM-CSF positive (*versus* GM-CSF negative) and memory (*versus* naïve) CD4 T cells, we scanned the consensus peaks for footprints and subsequently we scanned the identified footprints for TF binding motifs. For motif scanning we used the TRANSFAC database [47] that contains experimentally validated binding sites, consensus binding sequences (positional weight matrices) and regulated genes of eukaryotic TFs. Confirming the methodology used to identify footprints, they were enriched in differentially accessible regions as expected for the corresponding cell type comparisons (Supplementary Figure S3). Next, we defined the TFs whose binding sites were most enriched in peaks of GM-CSF positive or memory cells respectively (ranked based on differentially accessibility), and identified about 20 TFs each that passed the significance threshold (FDR<0.05) for enrichment (Figure 7A). These lists contained several TFs with a well-known role in T cells, such as SATB1, YY1, ETS- and EGR-family TFs, among other factors with a less defined role. Notably there was little overlap between the key TFs in GM-CSF positive and memory cells, suggesting that our strategy may have identified key factors to specifically define the GM-CSF positive T cell phenotype. The majority of these TFs were highly expressed on RNA level (Figure 7B), falling within the class of highly expressed genes (HEGs) that were suggested to be more functional than lowly expressed genes [50]. Interestingly however, most of these TFs were not differentially expressed themselves in the cell population comparisons under study (Figure 7C), suggesting that they may be regulated on post-transcriptional levels such as protein phosphorylation and intracellular localization which is well-known for many TFs. Hence, with a strategy exploring solely the transcriptome (or even proteome) without integrating ATAC-seq data, several key TFs would likely be missed, while our integrative strategy combining RNA-seq and ATAC-seq data successfully identified such TFs from limited amounts of primary human T cells.

**Figure 7.**
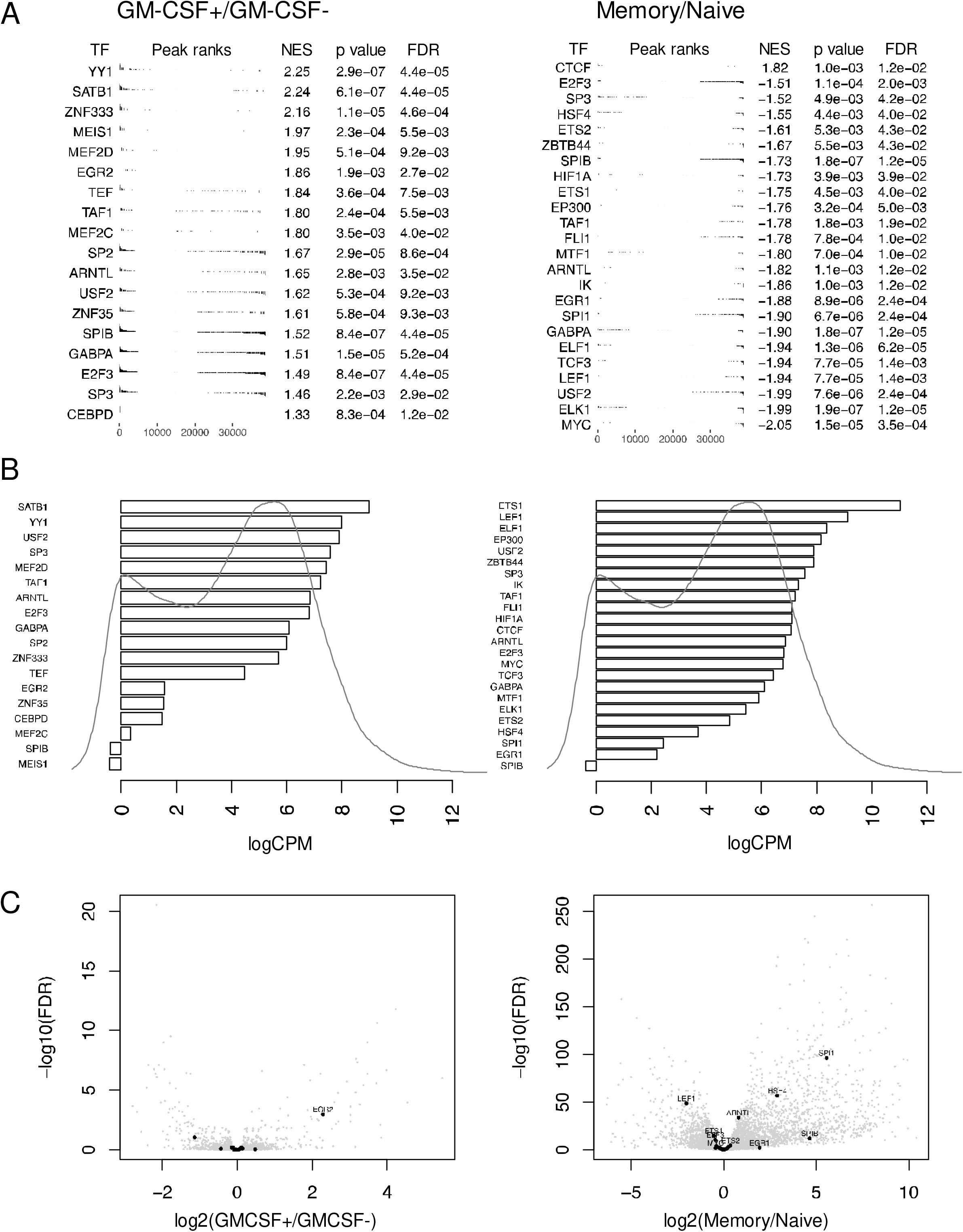
Identification of key TFs with global regulatory effect. (**A**) Rank based enrichment analysis of TF binding motifs in footprints within peaks that were ranked based on differential accessibility using the −log_10_(FDR) × sign(log_2_(fold change)) function. The analysis was performed separately for the GM-CSF+/GM-CSF- and Memory/Naïve cell comparison (left and right respectively). NES: normalized enrichment score. (**B**) Average expression levels of identified key TFs for both cell comparisons are shown as log(Count Per Million reads). To indicate the relative expression level of these TFs (bars) in view of the spectrum of lowly and highly expressed genes, the scaled probability density function is shown based on the distribution of the average expression of all genes. (**C**) Volcano plot shows the differential expression (effect size *vs.* significance) of the TFs identified as key regulators in each of the cell type comparisons (GM-CSF+/GM-CSF- and Memory/Naïve cell comparison; left and right). Key TFs are indicated as black dots, and those which are significantly differentially expressed (FDR<0.05) are labelled with their gene symbol.

### Integration of RNA-seq and ATAC-seq data identifies gene regulatory networks of GM-CSF positive and memory CD4 T cells

Having identified key TFs in GM-CSF positive and memory CD4 T cells, we were interested whether these factors regulated certain groups of target genes and whether several TFs may act together in a concerted fashion. Some TFs regulated a large number of target genes, and clusters of TFs were grouping together (Figure 8A, B). Exploring the co-binding between TFs in more detail showed that certain groups of TFs bound together to the same targets (Figure 8C, D). These included the TCF3:LEF1 pair that was always co-binding the same regions, as evident from the Memory/Naïve CD4 T cell contrast (Figure 8B, D). TCF/LEF family proteins act downstream of the Wnt pathway and it is well-known that they often display overlapping expression patterns and functional redundancy [65]. In accordance with our data on human memory and naïve CD4 T cells, binding motifs for TCF family members were recently also determined to be depleted in murine memory CD8 T cells as well as human memory CD4 T cells using ATAC-seq [35,39].

**Figure 8.**
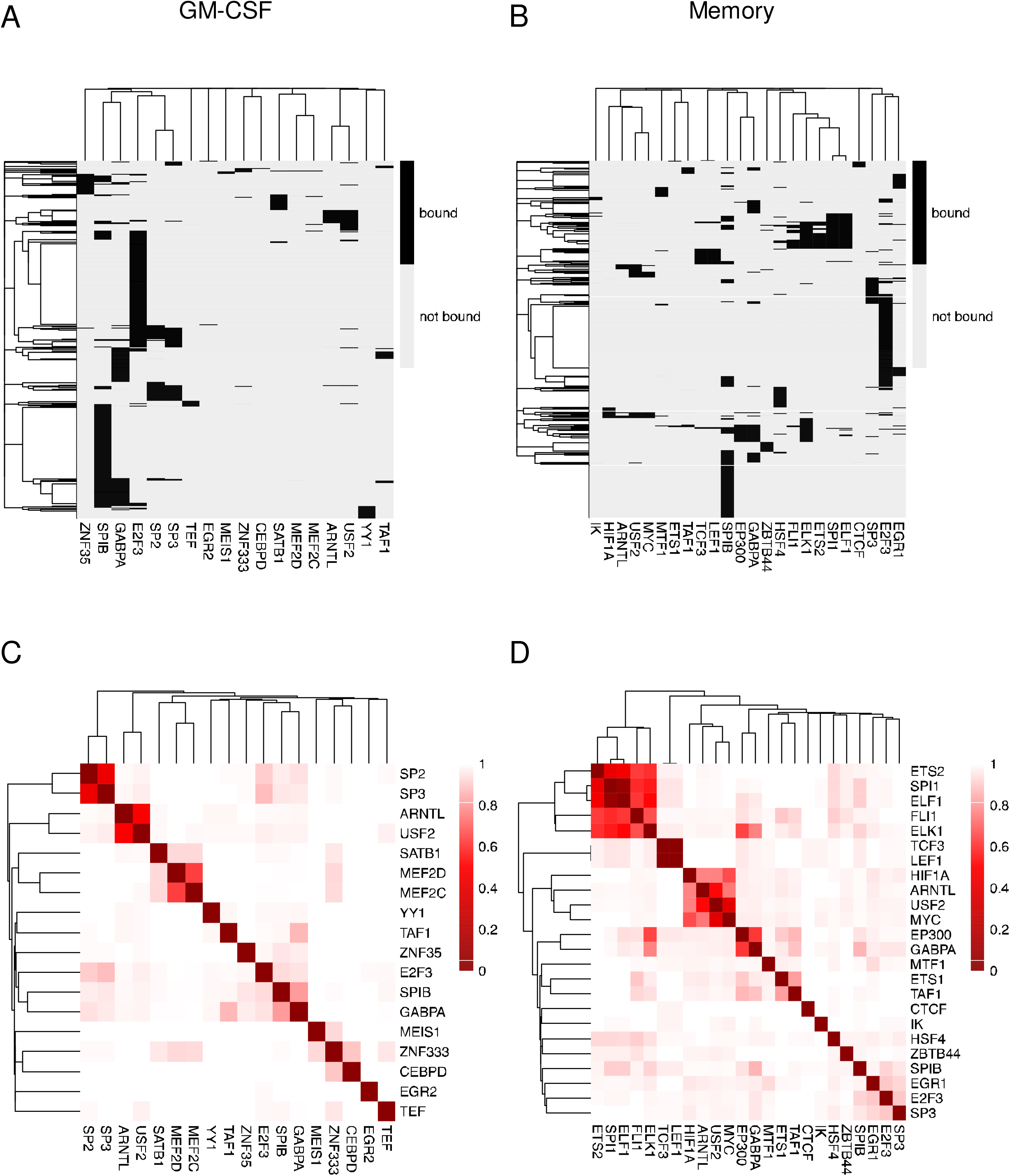
Network structure identifying shared TF binders across peaks and shared targets of key TFs. (**A, B**) Key TFs were identified as in Figure 7. The binary heatmap shows the binding of key TFs (columns) in footprints with ATAC-seq peaks (rows) that are either differentially accessible and/or that are assigned to a differentially expressed gene in the GM-CSF+/GM-CSF- cell comparison (A) or in the memory/naïve cell comparison (B). Black indicates that the key TF binds to the respective region. Rows and columns are clustered using binary distance and complete linkage. (**C, D**) Heatmap representation of the binary distance matrix of the key TFs using the binary vector of binding (“bound” or “not bound”) for distance calculation. The red color scale represents the distance; smaller distance represents more co-binding. Rows and columns are clustered using Euclidean distance and complete linkage. The analysis was performed for the GM-CSF+/GM-CSF- cell comparison (C) or the memory/naïve cell comparison (D).

Finally, by connecting the TFs to the genes assigned to their target regions, we generated a directed gene regulatory network representing GM-CSF positive (*versus* negative) and memory (*versus* naïve) CD4 T cells respectively (Figure 9). The source nodes were represented by the key TFs as selected above (key TFs from Figure 7A, B) and target nodes were selected when the region was either differentially accessible and/or the mRNA of the assigned gene was differentially expressed in the respective cell population comparisons. Both the GM-CSF as well as the memory comparisons led to similarly sized networks however, only some of the target and source nodes were shared between both networks (Figure 9A, B).

**Figure 9.**
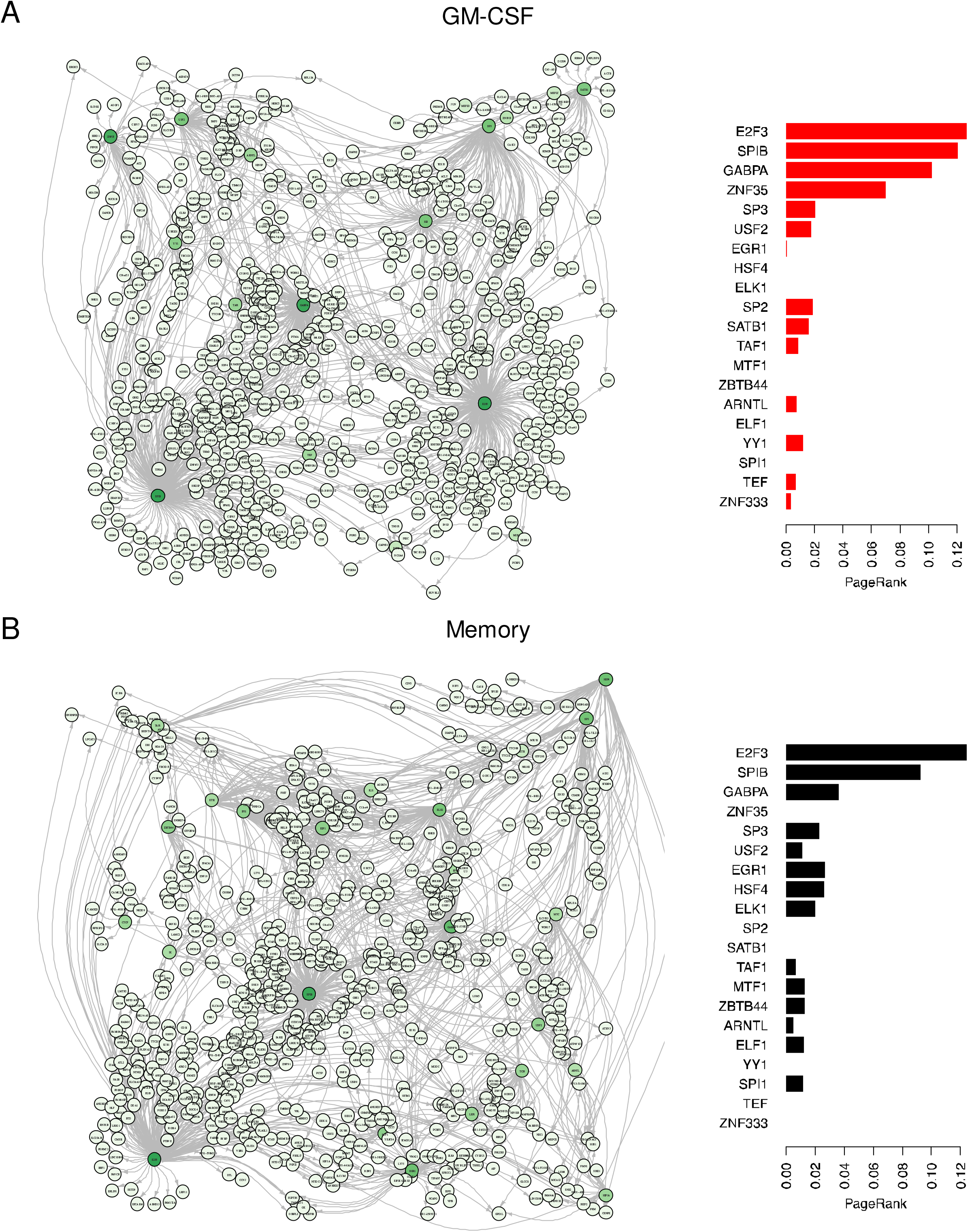
Gene regulatory network of GM-CSF+/GM-CSF- or Memory/Naïve CD4 T cells. (**A**) Gene regulatory network reflecting the signatures of GM-CSF positive *versus* GM-CSF negative CD4 T cells. Left panel: A directed network representing binding from source TFs to target peaks assigned to the indicated gene is shown. Source nodes were selected as described (Figure 7, key TFs) and target nodes were selected to be either differentially accessible (as a peak assigned to the respective node gene) and/or to be differentially expressed on mRNA level in the GM-CSF+/GM-CSF- contrast. To calculate the PageRank [51] as a measure of importance of the TFs (with TFs influencing more genes being more important), the edges were first inverted (not displayed), and the resulting PageRank value is represented by the color scale (light to dark green for lower to higher PageRank). Right panel: Those TFs having a PageRank value higher than the 99-percentile in either the GM-CSF and/or the Memory cell network (B) and their corresponding PageRank value in the GM-CSF network are displayed. (**B**): Same as (A), but for the Memory/Naïve CD4 T cell contrast.

To quantify the importance of the source nodes in each individual network as well as in the comparison of both networks, we calculated scores based on the PageRank algorithm [51]. To do such a calculation on a network where the outgoing edges from a TF reflect its importance, rather than the incoming edges like in the original application of the algorithm, the edges in our networks were inverted (solely for the purpose of calculating the PageRank values). The obtained PageRank values give information on the importance of a node based on the number of other nodes it influences (directly or indirectly). Some of the key TFs that were shared between both networks also had a relatively high PageRank in both networks, such as the TFs E2F3, SPIB and GABPA (Figure 9A, B), and may be general regulators of T cell activation/differentiation. Importantly, we also identified key TFs with a relatively high PageRank in the GM-CSF T cell regulatory network whose PageRank value in the memory/naïve network was 0. These TFs, namely ZNF35, SP2, SATB1, YY1, TEF and ZNF333 may, thus, be novel regulators specifically controlling the molecular signatures of GM-CSF positive CD4 T cells (Figure 9A, B).

## CONCLUSIONS AND OUTLOOK

In summary, we here provide interpreted high-quality novel RNA-seq and ATAC-seq data from primary human CD4 T cells. We provide a network of key TFs and their targets representing GM-CSF positive cells and memory CD4 T cells respectively. To our knowledge, this is the first study providing signatures of GM-CSF secreting cells isolated by capture assay without restimulation, which are likely as similar as possible to the state *in vivo* in humans. Very recently, others have performed a similar approach, capturing IFN-γ and IL-17 producing cells from human donors, notably also from MS patients [66], and the authors obtained important results on the transcriptional signatures of these cells, including genes distinguishing clinically stable *versus* active MS patients. This raises interesting prospects for studying GM-CSF positive cells specifically from MS patients as well by the methods and comparing to the data resource provided here. This will have important clinical implications given the likely important contribution of GM-CSF positive CD4 T cells to MS. Hu *et al.* [66] studied only a limited set of 418 genes in the captured IFN-γ and IL-17 secreting cells, while the methods employed in the present work enable genome-wide studies of DNA accessibility and RNA expression patterns. Notably, the authors [66] stated that their approach may “miss other disease related” genes and that “follow-up studies, such as RNA-sequencing analysis on TH17 and TH1/17 subsets isolated from MS patients, may help to identify more disease-related genes” which we propose should be extended to ATAC-seq and the GM-CSF producing Th subset as well. Future work should also address what distinguishes patterns and contributions of GM-CSF *versus* IFN-γ and IL-17 producing T cells particularly in MS patients. Although not from MS patients, another recent report [67] provides RNA-seq data of such captured cells from healthy donors however, PMA and ionomycin restimulation (which by itself drastically alters the T cell transcriptome) render it difficult to directly compare these data to ours. Interestingly, recent ATAC-seq data from murine GM-CSF *versus* IFN-γ and IL-17 producing T cells upon *in vitro* stimulation suggests that while all subsets showed distinct epigenetic profiles, GM-CSF producing cells were more related to IFN-γ than IL-17 producing T cells [10]. With an elegant fate-mapping system, that study [10] could also demonstrate that murine GM-CSF producing murine T cells display a stable epigenetic imprint. While such fate mapping systems are not possible with human cells, and also, capturing multiple cytokine-producing Th subsets in parallel from the same donor without artificially raising their fraction by PMA and ionomycin was not feasible in our study due to the limited cell numbers, our study provides data from unmanipulated *ex vivo* isolated human cells that may be most relevant for human diseases. Together, our data on naïve, memory, and GM-CSF positive CD4 T cells (and corresponding controls) can be exploited for a multitude of future studies for basic and translational immunology concerning autoimmune diseases. Beyond that, the results may have important implications for other diseases with involvement of GM-CSF producing cells, such as sepsis and COVID-19. Further, our data can be a test-bed for bioinformatics method development for the integration of RNA-seq and ATAC-seq data from the same samples, which may help understanding basic principles of gene regulation in primary eukaryotic cells.

## DATA AVAILABILITY

The data sets are available in the GEO repository under the SuperSeries accession number GSE119734 (with the SubSeries GSE119731 and GSE119732).

## AUTHOR CONTRIBUTIONS

S.E. and A.S. designed the project, designed and carried out all the experiments, interpreted results, prepared figures and wrote the paper; S.E. designed the computational data analysis pipeline and performed all the data analysis including visualization; D.G.C provided advice for data analysis and feedback on the manuscript; J.T. supervised research and provided feedback on the manuscript.

## ACKNOWLEDGEMENTS

The authors thank Matilda Eriksson and Peri Noori for performing RNA extractions, library preparation and quality control for RNA-seq, and next-generation sequencing, as well as for excellent general lab management; Sunjay Jude Fernandes for reagents and advice for the ATAC-seq protocol; and Gilad Silberberg for helpful discussions about footprinting and motif scanning (all from Computational Medicine Unit, Karolinska Institute). We thank John Andersson (Translational Immunology Unit, Karolinska Institute) for helpful suggestions and discussions.

## SUPPLEMENTARY MATERIALS

Supplementary Materials contain the following Figures and Tables: Supplementary Table S1A: Disease enrichment analysis based on gene lists ranked by gene expression in GM-CSF positive CD4 T cells (*versus* GM-CSF negative CD4 T cells). Supplementary Table S1B: Disease enrichment analysis based on gene lists ranked by gene expression in memory (*versus* naïve) CD4 T cells. Supplementary Table S2A: KEGG pathway enrichment analysis of genes assigned to consensus peaks. Supplementary Table S2B: GWAS catalog disease enrichment analysis of genes assigned to consensus peaks. Supplementary Figure S1: Purity of cell populations used in this study. Supplementary Figure S2: Expression profile and DNA accessibility signatures of human memory CD4 T cells. Supplementary Figure S3. Enrichment of footprints across peaks ranked based on differential accessibility.

---

**Supplementary Figure S1:**
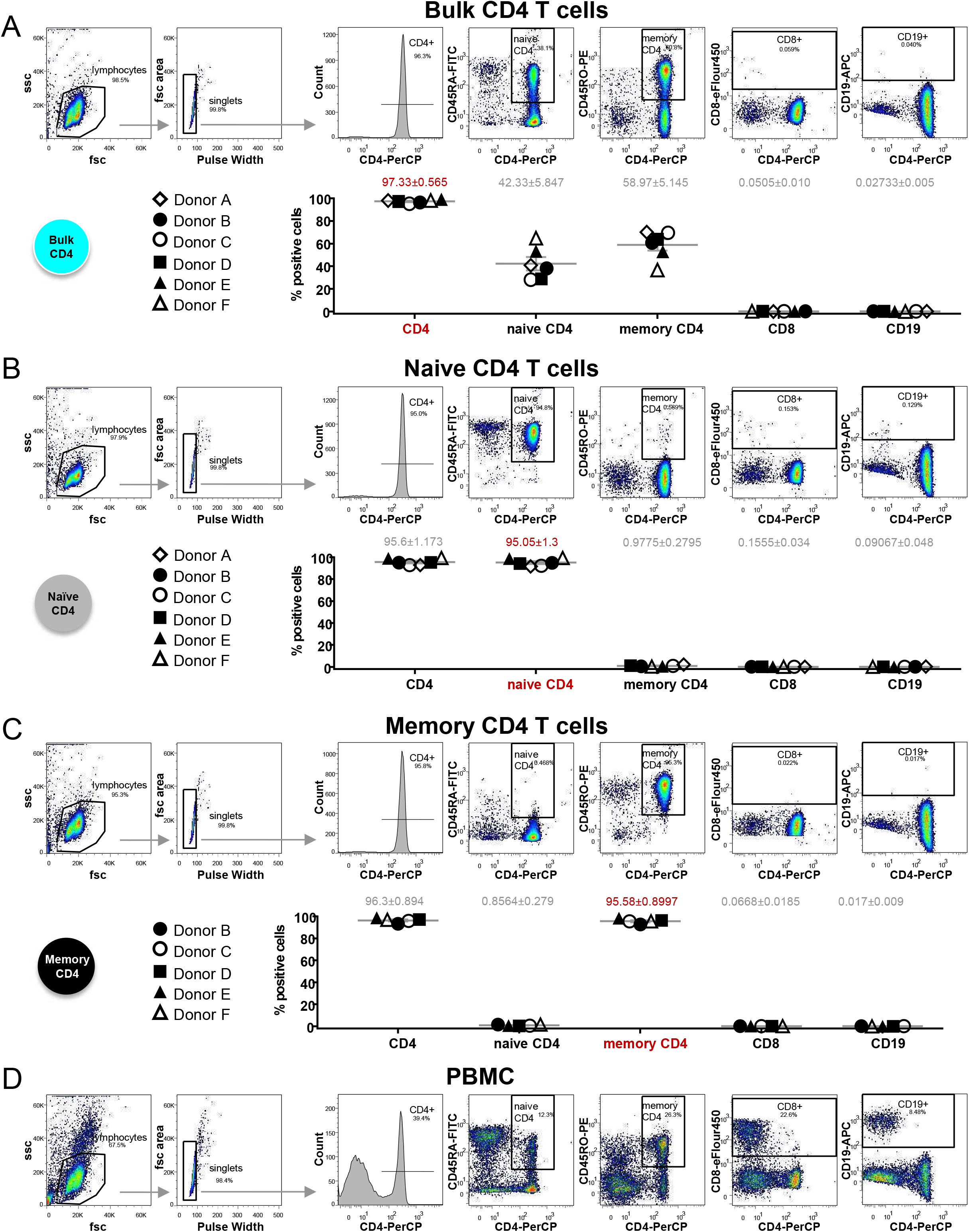
Purity of cell populations. Total CD4 T cells (**A**), naive CD4 T cells (**B**) and memory CD4 T cells (**C**) were isolated in parallel by negative magnetic isolation from human PBMCs. Cells were stained for the indicated surface markers and analyzed by flow cytometry. Upper panels show flow cytometry raw data for a representative donor each. The two scatterplots to the left show the pre-gating strategy; the 5 flow cytometry plots to the right are pre-gated on doublet-excluded lymphocytes. Gates to determine percentages of CD4+ cells, naive CD4+ cells, memory CD4+ cells, CD8+ cells and CD19+ cells are indicated. Lower panels show the summary data for the gates defined as above, for n=5 to 6 donors from 4 independent experiments. Same symbols (filled or unfilled) indicate donors processed within the same experiment. Horizontal grey lines represent mean values, and error bars in grey represent SEM. Mean±SEM values are indicated in grey, or highlighted in red font for the most relevant gate for the given isolated cell population. (**D**) shows PBMCs from a representative donor, which were always included in the staining as positive control. Mean±SEM values for PBMCs (n=6 donors) were as follows. CD4+ cell gate: 45.60±3.4%; naive CD4+ cell gate: 19.75±3.6%; memory CD4+ cell gate: 25.53±2.98%; CD8+ cell gate: 22.15±2.44 %; CD19+ cell gate: 7.91±0.64%.

**Supplementary Figure S2:**
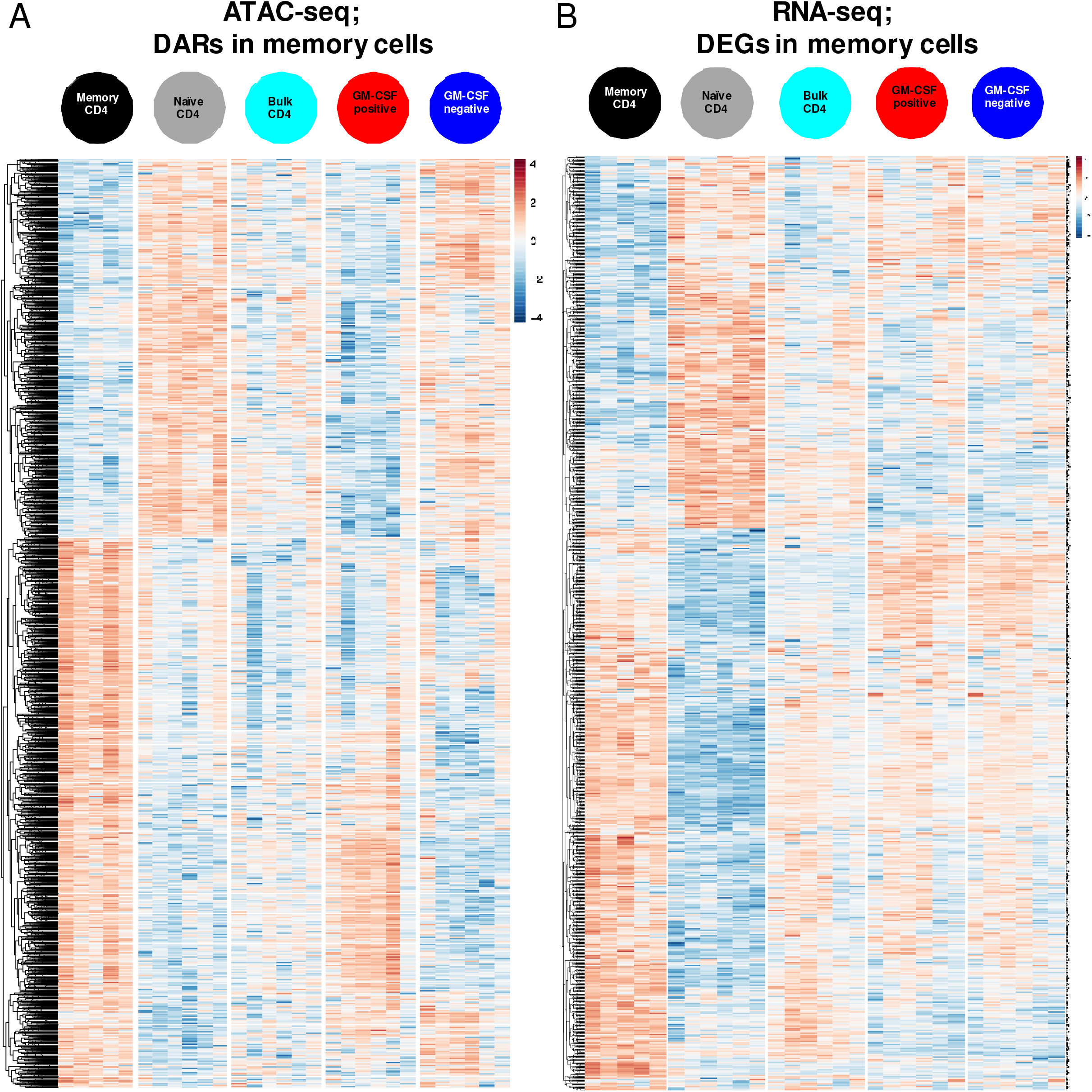
(**A**) Differentially accessible DNA regions in memory *versus* naive CD4 T cells (FDR<0.05 in this comparison) are plotted as a heatmap for their accessibility in all the five cell populations studied. log2(RPKM+1) is represented by the color scale, indicating accessibility (blue: low, red: high). Data were row-scaled and -clustered (Euclidean distance complete linkage). (**B**) Differentially expressed genes (RNA) in memory *versus* naive CD4 T cells (FDR<0.05 in this comparison) are plotted as a heatmap for their expression in all the five cell populations studied. Gene expression is displayed as log2(FPKM+1) with blue indicating low and red indicating high expression according to the color scale. Data were row-scaled and -clustered (Euclidean distance complete linkage).

**Supplementary Figure S3.**
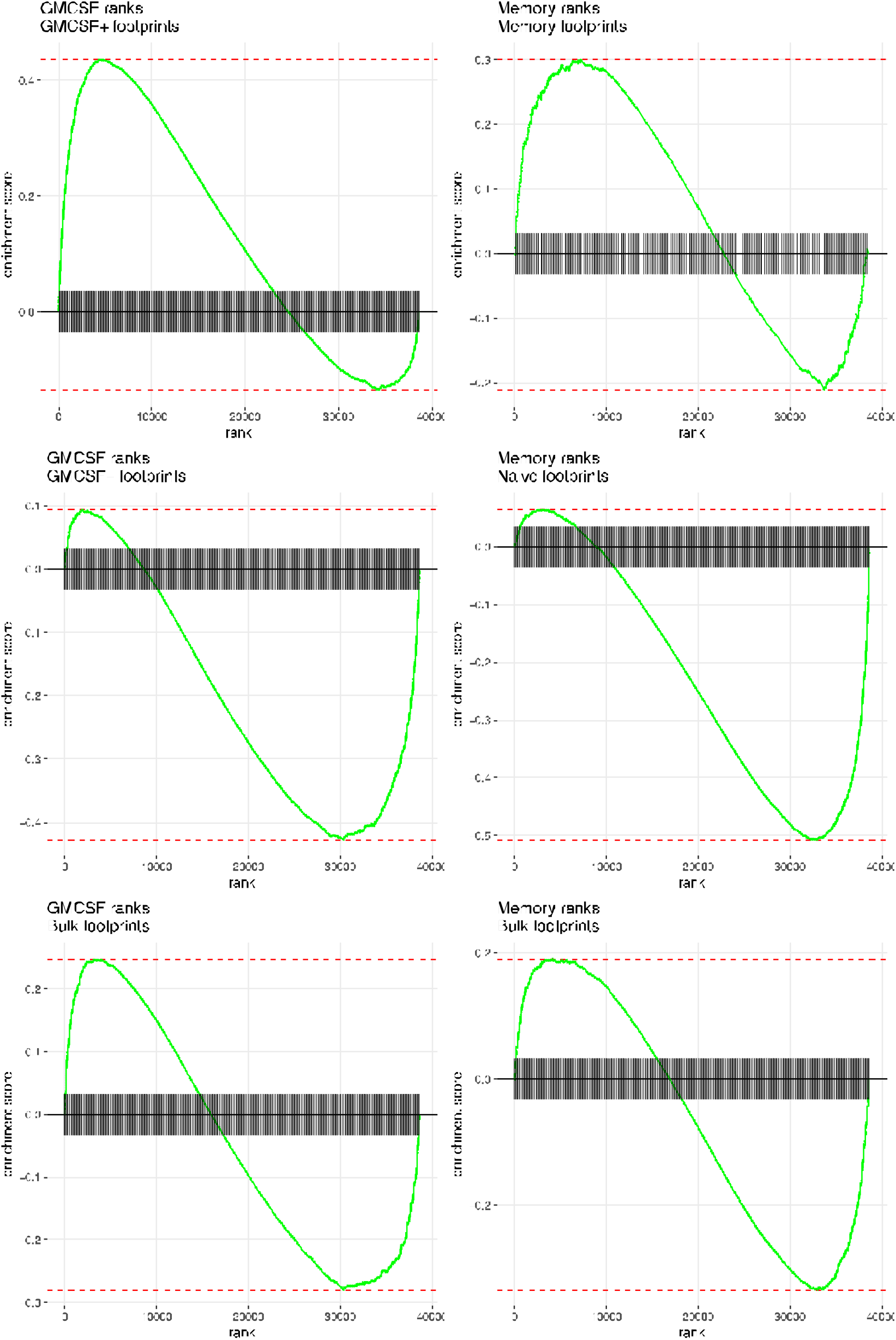
Enrichment of footprints across peaks ranked based on differential accessibility. Peaks were ranked based on their differential accessibility in the indicated contrast using the −log10(FDR) × sign(log2(Fold Change)). Peaks are represented on the horizontal axis and the enrichment score of the set of footprints identified in the indicated cell population is represented on the vertical axis.

